# SCA-1 micro-heterogeneity in the fate decision of dystrophic fibro/adipogenic progenitors

**DOI:** 10.1101/2020.05.14.096438

**Authors:** Giulio Giuliani, Simone Vumbaca, Claudia Fuoco, Cesare Gargioli, Ezio Giorda, Giorgia Massacci, Alessandro Palma, Alessio Reggio, Federica Riccio, Maria Vinci, Luisa Castagnoli, Gianni Cesareni

## Abstract

The term micro-heterogeneity refers to non-genetic cell to cell variability observed in a bell-shaped distribution of the expression of a trait within a population. The contribution of micro-heterogeneity to physiology and pathology remains largely uncharacterised. To address such an issue, we investigated the impact of heterogeneity in skeletal muscle fibro/adipogenic progenitors (FAPs) isolated from an animal model of Duchenne muscular dystrophy (DMD), the *mdx* mouse. FAPs play an essential role in muscle homeostasis. However, in pathological conditions or ageing, they are the source of intramuscular infiltrations of fibrotic or adipose tissue. By applying a multiplex flow cytometry assay, we characterised and purified from *mdx* muscles two FAP cell states expressing different levels of SCA-1. The two cell states are morphologically identical and repopulate each other after several growth cycles. However, they differ in their *in vitro* behaviour. Cells expressing higher levels of SCA-1 (SCA1-High-FAPs) differentiate more readily into adipocytes while, when exposed to a fibrogenic stimulation, increase the expression of *COL1A1* mRNA. In addition, SCA1-High-FAPs proliferate more extensively *ex vivo* and display more proliferating cells in dystrophic muscles. Adipogenesis of FAP cell states is inhibited *in vitro* by leukocytes from young dystrophic mice, while leukocytes isolated from aged dystrophic mice are less effective in limiting the adipogenesis of SCA1-High-FAPs suggesting a differential regulatory effect of the microenvironment on micro-heterogeneity. Our data suggest that FAP micro-heterogeneity is modulated in pathological conditions and that this heterogeneity in turn may impact on the behaviour of mesenchymal cells in genetic diseases.

## Introduction

The human body is estimated to contain approximately 200 different cell types ^1^. However, this estimate relies on the definition of “cell type” and on the experimental tools used to investigate cell populations and to define cell types. Developments in single cell technologies have challenged this view revealing that cell populations often display a higher molecular heterogeneity than originally reported ^2–4^. This new perspective raises several fundamental questions regarding origin and the impact of cell phenotypic or molecular heterogeneity on physiological and pathological processes^5^. Two different types of cell heterogeneities may be distinguished, macro-heterogeneity and micro-heterogeneity. The former refers to the presence of discrete cell sub-populations, while the latter can be viewed as a spread in the distribution of the expression of a trait in an seemingly homogeneous cell population ^6^. Micro-heterogeneity generates metastable states that originate from a stochastic transcriptome-wide noise ^7^. This kind of cell to cell variability has been reported to modulate some important physiological mechanisms as, for instance, the differentiation decisions of a multipotent mouse hematopoietic cell line and the asymmetric DNA segregation during Muscle Satellite Cells (MuSCs) division ^7,8^. However, the involvement of micro-heterogeneity in disease is only poorly explored.

We asked whether the heterogeneity observed in mononuclear muscle cell progenitors has any impact on and is affected by myopathies such as Duchenne muscular dystrophy (DMD). DMD is an X-linked genetic disease caused by mutations in the dystrophin gene ^9^. Sarcolemma instability originating from this condition leads to chronic damage, progressive muscle wasting and deposition of ectopic tissues, such as intramuscular adipose tissue (IMAT) and fibrotic tissue ^10,11^. The source of ectopic tissues is a population of mesenchymal cells known as fibro/adipogenic progenitors (FAPs) ^12^. FAPs reside in the endomysium of skeletal muscle and are characterised by the expression of the surface proteins stem cell sntigen-1 (SCA-1 also known as lymphocyte antigen 6A-2/6E-1, Ly6a) and CD140a (also known as platelet derived growth factor receptor alpha, PDGFR-alpha) ^13–16^. Recently, single cell approaches have revealed heterogeneity in FAPs and have provided some evidence of its involvement in muscle development and dystrophy in mouse models ^17,18^.

Here, we characterise two dynamic cell states in FAPs from muscles of *mdx* mice and describe how this heterogeneity impacts their properties *ex vivo* and *in vivo*. We show that FAP cell states, characterised by different levels of SCA-1 expression, differ in differentiation and proliferation potential. Furthermore, we show that the muscle microenvironment differentially modulates the behaviour of the two FAP cell states by affecting the cell propensity to differentiate into adipocytes in an age-dependent manner. These results suggest that micro-heterogeneity takes part in fate decision of a mesenchymal cell population involved in the pathogenesis of a genetic disease.

## Results

### SCA-1 expression defines two FAP populations

We have recently shown that purified FAPs are a heterogeneous population including several sub-populations ^18^. Here we use a multiplex flow cytometry approach to characterise FAP heterogeneity from a suspension of mixed populations derived from skeletal muscles. We first isolated mononuclear cells from hind limb muscles of 6-8 weeks old wild type and *mdx* mice. In this time window, muscles of dystrophic mice are in a regeneration phase ^19,20^. Cells were stained with fluorescent antibodies against 9 surface antigens (CD45, CD31, Integrin alpha-7, SCA-1, CD140a, CD140b, CD146, CD34, CD90.2) to distinguish the main mononuclear cell populations residing in the skeletal muscle: leukocytes (CD45^+^), endothelial cells (CD45^-^ CD31^+^), muscle satellite cells (MuSCs) (CD45^-^ CD31^-^ Integrin alpha-7^+^) and FAPs (CD45^-^ CD31^-^ Integrin alpha-7^-^ SCA-1^+^). The antigen expression profiles of single cells were analysed by a mixed approach whereby an automatic clustering is followed by expert manual refinement (Figure 1A and details in supplementary materials). This approach yielded 4 cell clusters corresponding to the main muscle mononuclear cell populations (Figure 1B) and 11 unassigned clusters (Supplementary Figure 1D), which were characterised by an antigen profile that could not be assigned to any of the main mononuclear cell populations in the muscle niche. FAPs amount to 2.6% of the mononuclear cells in the wild type mouse, whereas their abundance increases to 5.7% in the *mdx* mouse (Supplementary Figure 1E).

**Figure 1.**
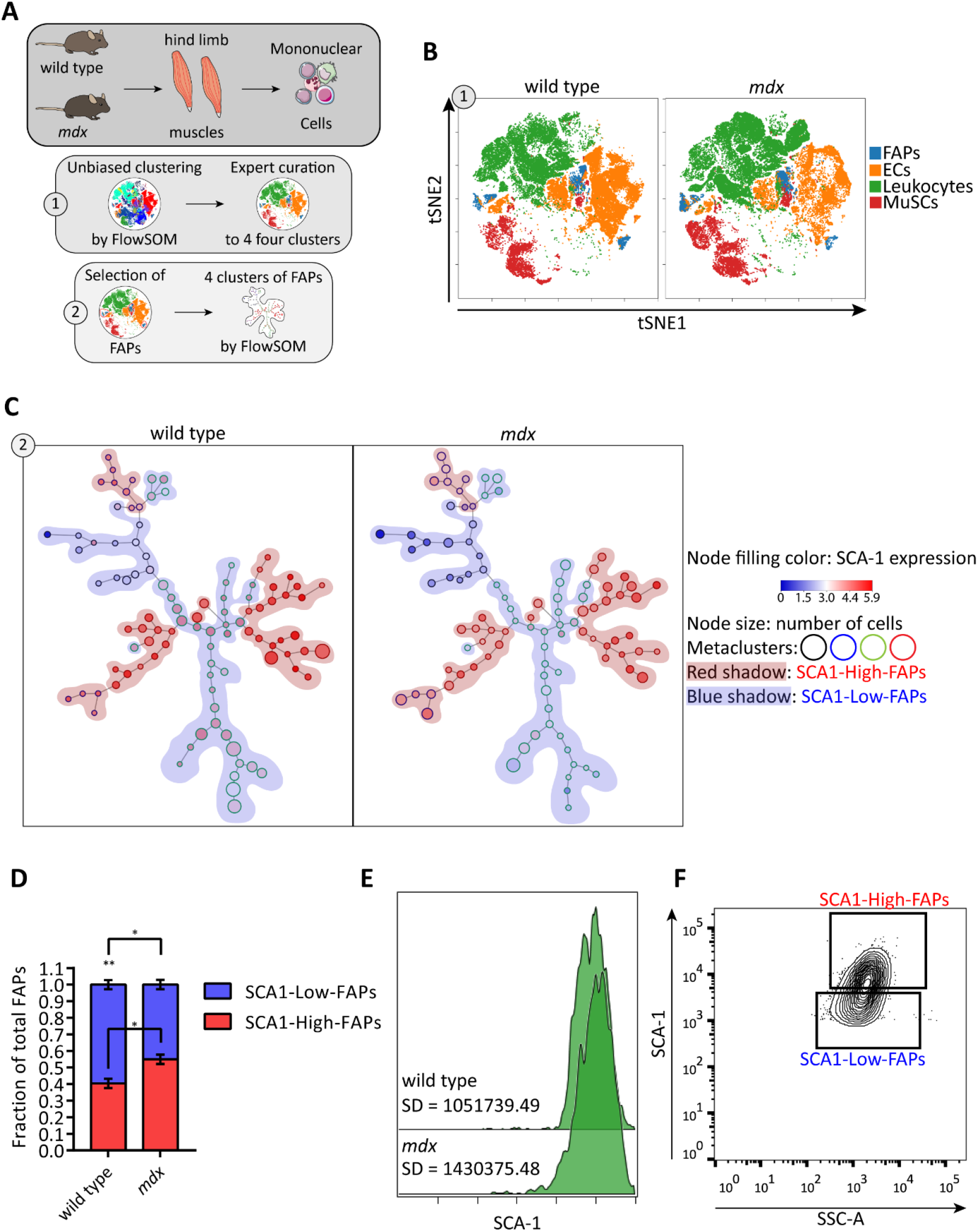
Micro-heterogeneity of SCA-1 in fibro/adipogenic progenitors. (A) Schematic representation of the workflow applied to study FAP heterogeneity. (B) Representative viSNE maps of total mononuclear cells isolated from young wild type and *mdx* mice. The four clusters produce by FlowSOM algorithm were mapped onto the viSNE maps and correspond to the following mononuclear cell populations: FAPs (blue), endothelial cells (orange), leukocytes (green) and muscle satellite cells (MuSCs, red). (C) Representative Self Organizing Maps (SOMs) of FAPs identified in (A). SOMs were obtained with the FlowSOM algorithm. Each node represents a cluster of cells, nodes with similar expression profile are linked by an edge. Colour of nodes indicate SCA-1 expression level. Node outlines indicate the four metaclusters (red, blue, black and green) obtained by the algorithm. Red and blue shadings highlight FAPs expressing high levels (red and blue metaclusters, called SCA1-High-FAPs) and low levels (black and green metaclusters, called SCA1-Low-FAPs) of SCA-1. (D) Stacked bar plot showing the fraction of SCA1-High-FAPs and SCA1-Low-FAPs in wild type and *mdx* mice. Data are presented as mean ± SEM. Statistical significance was estimated by a One-way ANOVA, * p ≤ 0.05, ** p ≤ 0.01. n = 4. (E) Representative SCA-1 histograms of FAPs from wild type and *mdx* mice identified in (B) and their standard deviations (SD) showing a typical micro-heterogeneity profile. (F) Sorting strategy to decompose SCA-1 micro-heterogeneity and to isolate SCA1-High-FAPs and SCA1-Low-FAPs from *mdx* mice. Complete strategy, Fluorescence minus one (FMO) controls and cell states purity in Supplementary Figure 3.

We next focused on FAPs and we chose to study the expression variability of 6 antigens (SCA-1, CD140a, CD140b, CD146, CD34, CD90.2) associated to the FAP immunophenotype ^13,14,18^. By applying the same clustering strategy used for the total mononuclear populations, we obtained 4 metaclusters represented by red, blue, green and black outlined nodes of the Self Organizing Maps (SOMs) produced by the FlowSOM algorithm (Supplementary Figure 2 and Figure 1C). Interestingly, we noticed metaclusters that express high level of SCA-1 (red and blue outlined nodes highlighted by red shading in Figure 1C) and metaclusters expressing low level of SCA-1 (black and green outlined nodes highlighted by blue shading in Figure 1C). These two FAP populations are equally represented in the *mdx* mouse, while FAPs expressing low level of SCA-1 are predominant in the wild type model (Figure 1D). The intensity distribution of SCA-1 shows a single peak and a large standard deviation (Figure 1E), features that characterise the cell to cell variability dubbed micro-heterogeneity ^6^.

### SCA1-High-FAPs and SCA1-Low-FAPs are cell states in dynamic equilibrium

The observation that FAPs display micro-heterogeneity prompted us to design a sorting strategy to isolate, from the *mdx* mouse model, FAPs (CD45^-^ CD31^-^ Integrin alpha-7^-^ SCA-1^+^) expressing high and low levels of SCA-1, henceforth called SCA1-High-FAPs and SCA1-Low-FAPs respectively (Figure 1F; refer to Supplementary Figure 3A for the complete gating strategy, to the Supplementary Figure 3B-C for controls and population purity). Similar sorting strategies have already been used to resolve micro-heterogeneity ^7,8,21^. Sorted FAPs are significantly enriched for CD140a, as assessed by immunofluorescence (Supplementary figure 3D) and by flow cytometry (Supplementary Figure 3E).

Next, we asked whether FAPs expressing high and low levels of SCA-1 represent stable sub-populations or rather dynamic cell states. Terminology in the cell plasticity field is still debated ^22^. Here we define a “sub-population” of a given cell type as a group of cells identified by a discrete expression of specific markers and capable of maintaining identity when grown for several duplications *ex vivo*. Instead, considered a seemingly homogeneous cell population, we talk of a “cell state” when referring to a group of cells with an expression profile skewed when compared to the symmetrical expression distribution of the parent population. Cell states are in a dynamic equilibrium therefore they lose their identity after *in vitro* expansion ^7^.

To address this issue, we performed a re-population experiment in which SCA1-High-FAPs and SCA1-Low-FAPs were allowed to proliferate *ex vivo*. After 9 days, we analysed SCA-1 expression by flow cytometry. The SCA-1 intensity distributions of freshly isolated cells are clearly separated, whereas in expanded cells they overlap and restore the expression profile of the whole FAP compartment (Figure 2A, gating strategies and control in Supplementary Figure 3A-E). This observation suggested that SCA1-High and SCA1-Low-FAPs are dynamic states capable of repopulating each other.

**Figure 2.**
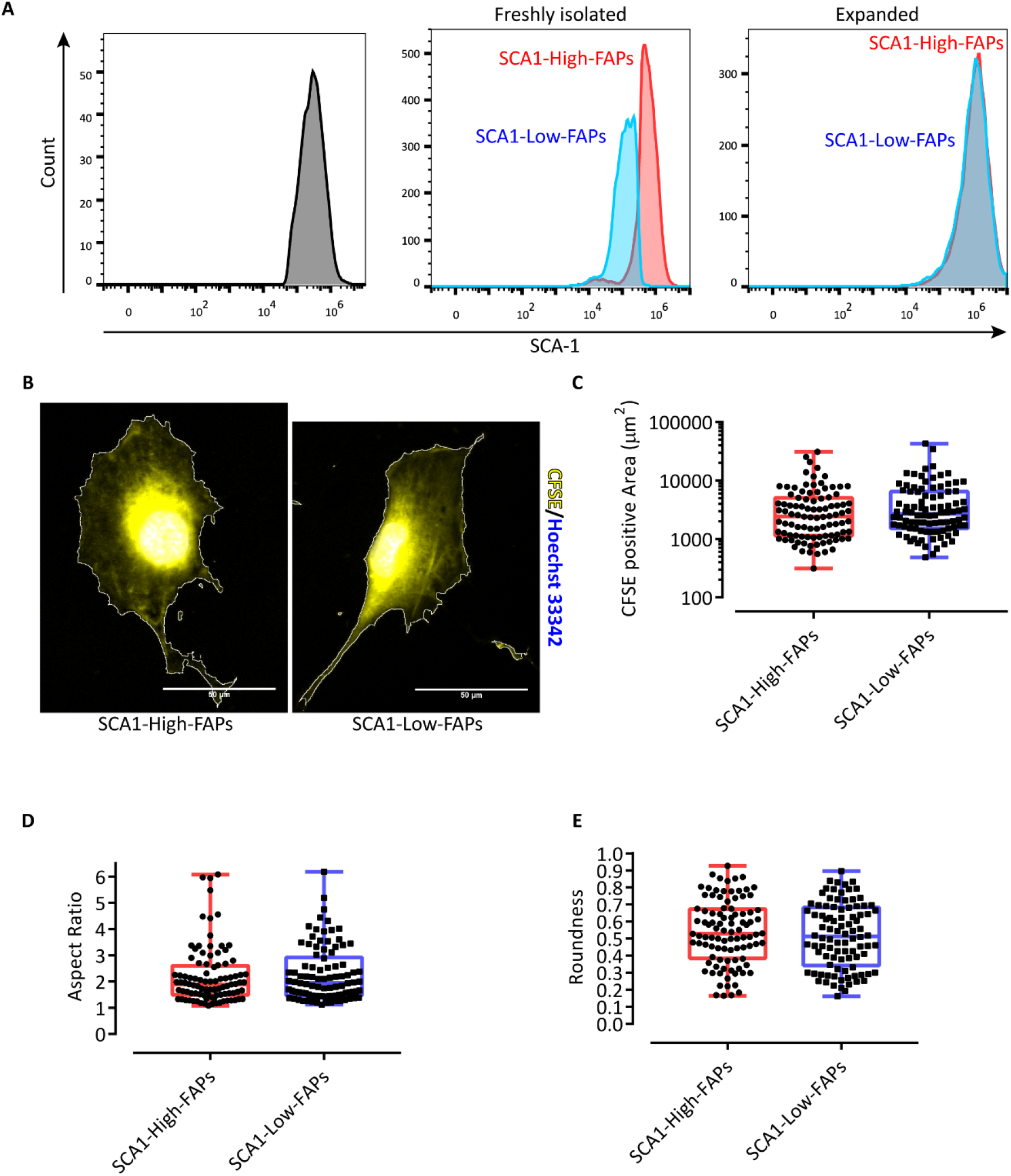
SCA1-High-FAPs and SCA1-Low-FAPs from *mdx* mouse are two cell states. (A) SCA-1 histograms of the whole FAP compartment (left), freshly isolated cell states (middle) and cell states expanded 9 days in Cytogrow (right). FAP compartment was analysed as followed: CD45^-^ CD31^-^ cells isolated by MACS were stained with antibodies against Integrin alpha-7 and SCA-1; next a gating strategy (Supplemetary Figure 4B; unstained cells in Supplementary Figure 4A) was applied to obtain Integrin alpha-7^-^ SCA-1^+^ cells. Cell states were isolated as described and then analysed by flow cytometry (gating strategy in Supplementary Figure 4D and 4E for SCA1-High-FAPs and SCA1-Low-FAPs respectively; controls in Supplementary Figure 4C). n =3. (B) Representative crop of micrographs of SCA1-High-FAPs (left) and SCA1-Low-FAPs (right) cultured *ex vivo*. Cytoplasms were stained with CFSE (yellow) and nuclei were counterstained with Hoechst 33342 (blue). Scale bar 50 μm. (C), (D) and (E) Box plots representing CFSE positive area, aspect ratio (AR) and roundness of SCA1-High-FAPs (n = 96) and SCA1-Low-FAPs (n = 94) from three different biological replicates. Whiskers are minimum and maximum values of the distribution. Statistical significance was estimated by a two tailed Mann-Whitney test after a normality test.

We next investigated whether SCA1-High-FAPs and SCA1-Low-FAPs could be distinguished *in vitro* according to their morphology. To this end we stained their cytoplasms with CarboxyFluorescein Succinimidyl Ester (CFSE) and we selected three suitable descriptors to characterise cell shape ^23^. Roundness and aspect ratio (AR) are parameters that describe round cells and elongated cells respectively (Supplementary figure 4F). The third descriptor is cell size, which was estimated by the CFSE positive area. As a control we included in this analysis MuSCs since they acquire characteristic shapes during myogenic differentiation. Indeed, in the MuSC sample we are able to identify cells that exhibit high roundness/low AR and cells with low roundness/high AR (Supplementary figure 4G-J). The former round cells are proliferating myoblasts, the latter elongated cells are differentiating myocytes ^24^. Moreover, these myogenic cells are smaller than SCA1-High-FAPs or SCA1-Low-FAPs (Supplementary Figure 4H). Focusing on SCA1-High-FAPs and SCA1-Low-FAPs, we found that these cells are indistinguishable by any of the three shape describing parameters thereby excluding cell morphology or size as grounds for the observed heterogeneity (Figure 2B-E).

In conclusion, using as a criterion micro-heterogeneity in SCA-1 expression, we identified two FAP cell states that populate the skeletal muscle of the *mdx* mouse. The two cell states are occupied by morphologically identical cells and are dynamic as they readily lose their identity and acquire similar SCA-1 expression profiles when grown in culture.

### SCA1-High-FAPs have a higher adipogenic propensity than SCA1-Low-FAPs

The impact of macro-heterogeneity on the differentiation potential of adipose progenitors has been characterised in the adipose tissue ^25^. Whether micro-heterogeneity also plays a similar role in adipogenic progenitors in the skeletal muscle, and in differentiation in general, remains unexplored. The FAP adipogenic potential is restrained in the skeletal muscle by negative signals. In pathological conditions, however, the several cues controlling this process are deregulated leading to the deposition of IMAT ^18,26,27^. Muscles of the *mdx* mouse develop ectopic adipose tissue only in old animals ^27^. However, when cultured *ex vivo* in the absence of the anti-adipogenic signals of the muscle niche, freshly isolated FAPs from young *mdx* mice readily differentiate into adipocytes ^28^. We asked whether the micro-heterogeneity in the expression of SCA-1 has any impact on the *ex vivo* adipogenic potential.

The two cell states of FAPs purified from *mdx*-mice muscles were cultured for 5 days in growth medium (GM). After this short expansion period, adipogenic differentiation was induced by incubating cells with the adipogenic induction medium (AIM), GM supplemented with 1 μg/ml insulin, 1 μM dexamethasone and 0.5 mM of 3-isobutyl-1-methylxanthine (IBMX), for three days followed by additional two days in the adipogenic maintenance medium (MM), GM supplemented with 1 μg/ml insulin (Figure 3A). At the end of the differentiation process, we stained the lipid droplets using Oil Red O (ORO). We noticed little differentiation in both cell states when incubated in GM in the absence of adipogenic stimuli. Interestingly in cells incubated with the adipogenic medium (AM), the percentage of adipocytes in SCA1-High-FAPs is significantly higher than the one observed in SCA1-Low-FAPs (Figure 3B and 3D). This different behaviour may be the result either of a different propensity to undergo adipocyte commitment or to a delay in adipogenic differentiation. We addressed this point by evaluating the expression of peroxisome proliferator-activated receptor gamma (PPAR-gamma) by immunofluorescence. This transcription factor is a master regulator of adipogenesis and is not expressed in FAPs soon after isolation ^13,14^. PPAR-gamma expression is significantly higher in SCA1-High-FAPs (Figure 3C and 3D) suggesting that adipocyte commitment may be impaired in SCA1-Low-FAPs. To further support this conclusion, we allowed a longer time for FAPs to differentiate into mature adipocytes by extending the incubation in MM to 11 days (Figure 3E). This long-term insulin stimulation allows cells to display their full adipogenic potential. In this condition, both cell states differentiate more efficiently and adipocytes are more mature compared to what observed in the standard protocol (Figure 3G). However, also in this condition, a larger percentage of SCA1-High-FAPs yielded mature adipocytes (Figure 3F). At the end of the differentiation process SCA1-High-FAPs reach a higher confluence when compared to SCA1-Low-FAPs (Supplementary Figure 5A and 5B). To rule out the possibility that the observed differences in the adipogenic differentiation of FAP cell states could be a consequence of a different kinetics in reaching cell confluence, we plotted the total number of nuclei per field over the total number of adipocytes per field from the experiment in Figure 1A-D and we calculated the trend lines ^28^. Even though the trend line of SCA1-High-FAPs clearly reveals a correlation between cell onfluence and adipogenic differentiation, the trend of SCA1-Low-FAPs shows no increment of adipocytes at increasing nuclei number (Supplementary Figure 5C). This analysis is consistent with SCA1-Low-FAPS having lower adipogenic propensity, irrespective of confluence process.

**Figure 3.**
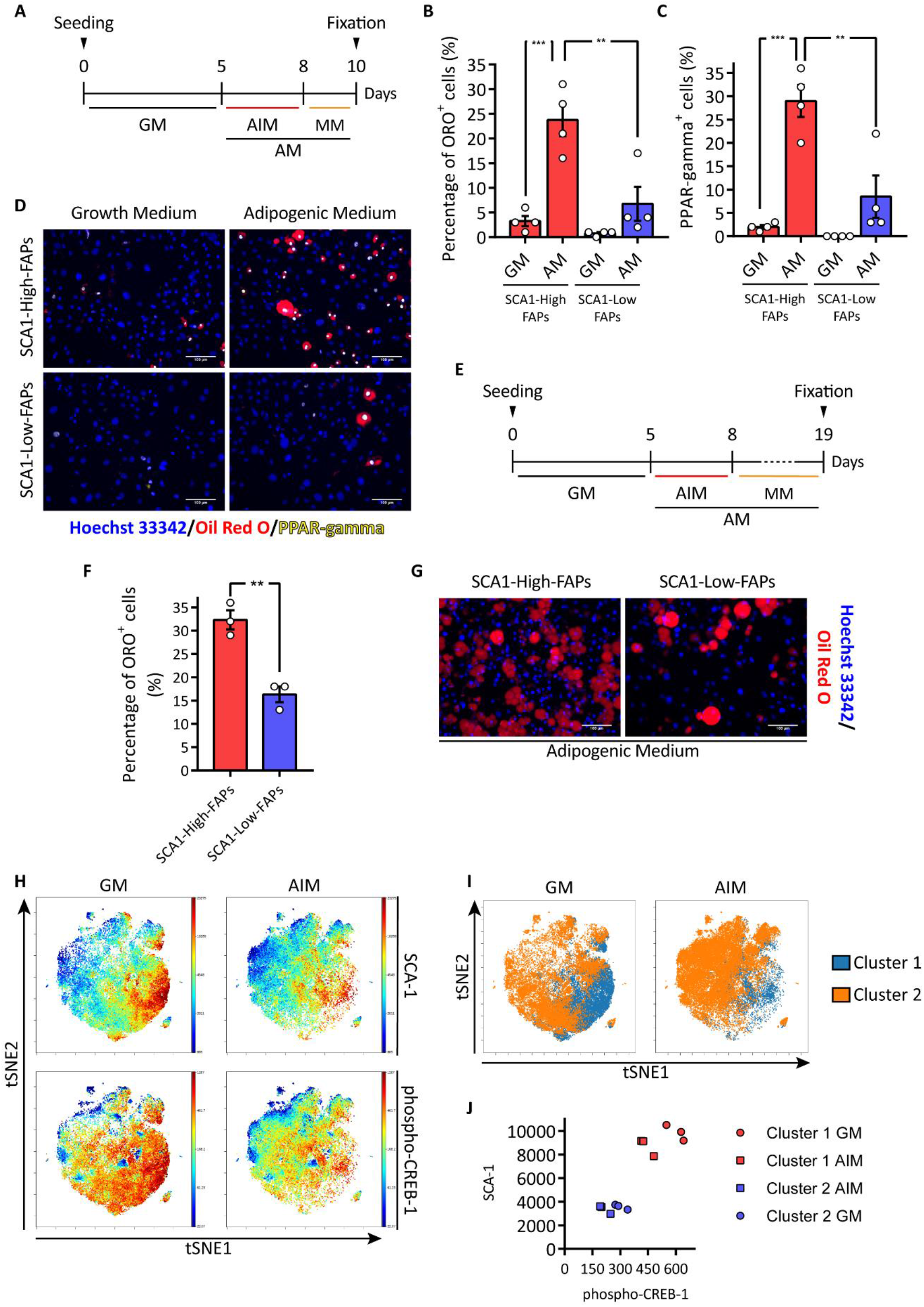
SCA1-High-FAPs differentiate more efficiently into adipocytes. (A) Experimental design to induce adipogenic differentiation of *mdx* FAP cell states. GM = growth medium; AIM = adipogenic induction medium; MM = maintenance medium. (B) and (C) Bar plots showing the percentage of ORO positive cells and PPAR-gamma positive cells per field. AM = adipogenic medium (AM = AIM + MM). n = 4. Statistical analysis was performed by a Two-way ANOVA. (D) Representative micrographs of (B) and (C). Cells were immunolabelled for PPAR-gamma (yellow) and nuclei were counterstained with Hoechst 33342 (blue). Lipid droplets were stained with ORO (red). (E) Experimental design applied to obtain fully differentiated adipocytes. (F) Bar plot indicating the percentage of ORO positive cells. n = 3. Statistical significance was calculated through a Student t-test. (G) Representative micrographs of (F). Lipid droplets in red (ORO staining) and nuclei in blue (Hoechst 33342). (H) Representative viSNE maps showing the expression of SCA-1 and phospho-CREB-1 assessed by mass cytometry in FAPs isolated from *mdx* mouse by MACS and cultured 72 hours either in GM or AIM. (I) Cluster 1 (in blue) and cluster 2 (in orange) obtained applying the FlowSOM algorithm and then mapped onto viSNE maps. (J) Plot representing the expression in arbitrary units of SCA-1 and phospho-CREB-1 identified in (J). (H) Representative viSNE maps of FAPs isolated by MACS from *mdx* mice and then incubated either to GM or to AIM. The upper maps show expression level of SCA-1, while the lower maps show the level of phospho-CREB-1. n = 3. (I) Representative viSNE maps showing the two clusters produced by the clustering approach. (J) Dot plot showing the expression level of SCA-1 and the level of phospho-CREB-1 (both in arbitrary units) in the metaclusters from (I). Data are presented as mean ± SEM. ** p ≤ 0.01, *** p ≤ 0.001. Scale bars 100 μm.

The correlation between SCA1 expression and adipogenic potential is also confirmed by a single cell approach. FAPs were isolated from muscles of *mdx* mice by MACS and then were incubated in either GM or AIM. After 72 hours single cells were analysed by mass cytometry using a panel of antibodies against 18 proteins (CD45, CD146, cleaved CASP-3, CD34, phospho-EGFR, CD140a, pRb, CD140b, Vimentin, CD90.2, phospho-STAT3, Integrin alpha-7, CXCR-4, SCA-1, CD31, c-Kit, Actin, phospho-CREB-1). viSNE maps highlight a correlation between phospho-CREB-1, a marker of early adipogenic differentiation, and SCA-1 expression in cells maintained in the growth medium (Figure 3H). By applying the FlowSOM algorithm, we observed two clusters expressing high levels of SCA-1 and phospho-CREB-1 (the blue cluster) and low levels of these proteins (the orange cluster) (Figure 3I-J). Consistently the levels of phospho-CREB-1 decrease in cells after three days of differentiation but remain higher in cells spontaneously differentiating in growth medium as CREB-1 phosphorylation is an early event in adipogenesis ^29,30^.

Collectively, these results indicate that the different SCA-1 expression levels in the FAP cell states correlate with a different adipogenic potential. SCA1-Low-FAPs are characterised by a reduced adipogenesis, whereas SCA1-High-FAPs engage in this differentiation path more readily.

### FAP cell states differ in the expression of fibrogenic genes

Abnormal deposition of extracellular matrix (ECM) is another pathogenic hallmark of muscular dystrophies. Differently from fat deposition this condition also develops in young dystrophic mice. Myofibroblasts differentiate from several cell types and, among these, FAPs are considered the main source of fibrotic scars ^12^. Upon differentiation, myofibroblasts synthetize alpha-smooth muscle actin (alpha-SMA) to form positive stress-fibres and increase the production of ECM proteins ^31^. Transforming growth factor beta-1 (TGF-beta-1) promotes fibrogenic differentiation of FAPs *in vitro* and takes part in fibrosis *in vivo* ^12,32^.

We asked whether FAP heterogeneity, in addition to modulating adipogenesis, had also an impact on fibrogenesis. After 5 days in growth medium (GM), FAPs were exposed for 5 additional days to a fibrogenic medium (FM) containing TGF-beta-1 (Figure 4A). Next, the expression of alpha-SMA was monitored by immunofluorescence. Both cell states respond to FM by increasing the alpha-SMA-positive area. The alpha-SMA positive area surrounding SCA1-Low-FAP cells was twice as large as that of SCA1-High-FAP cells (Figure 4B-C, nuclei count in Supplementary Figure 6). The SMA-positive area in samples exposed only to GM also tends to be higher in SCA1-Low-FAPs.

**Figure 4.**
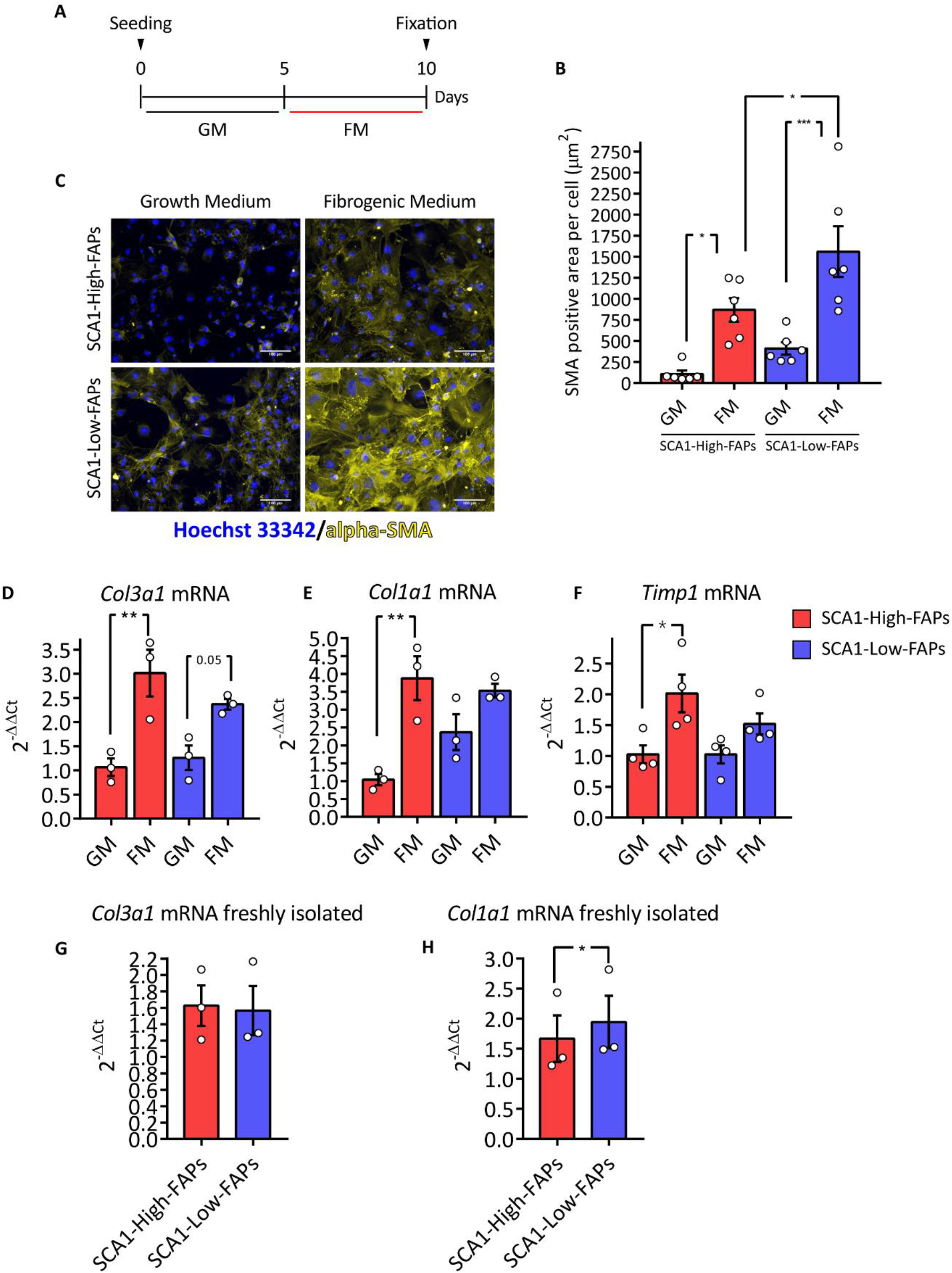
Fibrogenic differentiation of *mdx* FAP cell states. (A) Experimental design to induce fibrogenic differentiation of *mdx* FAP cell states. GM = growth medium; FM = fibrogenic medium. (B) Bar plot presenting alpha-SMA positive area per cell in each field. n = 6. Statistical significance was evaluated using Two-way ANOVA. (C) Representative microphotographs of (B). Immunofluorescence against alpha-SMA (yellow) and nuclei counterstaining (blue, Hoechst 33342). Scale bar 100 μm. (D), (E) and (F) Bar plots showing *Col3a1, Col1a1* and *Timp1* mRNA expression. Following 5 days of differentiation samples were analysed by Real-Time PCR using 2^-ΔΔCt^ comparative method. Tubulin was set as reference gene. n = 3. Statistical significance was estimated through a Two-way ANOVA. (G) and (H) Bar plots showing *Col3a1* and *Col1a1* mRNA expression immediately after sorting. Samples were analysed by Real-Time PCR using 2^-ΔΔCt^ comparative method. *Tubulin* mRNA was set as reference gene. n = 3. Statistical significance was evaluated by a Student t-test. Data are presented as mean ± SEM. * p ≤ 0.05, ** p ≤ 0.01, *** p ≤ 0.001.

We also monitored the mRNA levels of *Col3a1* and *Col1a1* genes upon fibrogenic induction. These collagen chains are components of the skeletal muscle ECM and are produced by FAPs upon TGF-beta-1 treatment *in vitro* ^12^. Both cell states respond to FM by increasing the expression of *Col3a1* mRNA (Figure 4D). However, only SCA1-High-FAPs significantly increased the expression of *Col1a1* mRNA upon induction of differentiation (Figure 4E). By looking at the mRNA levels in the absence of induction of fibrogenesis (GM), SCA1-Low-FAPs tend to express higher level of *Col1a1*. This difference was significant in freshly isolated cells (Figure 4H). We found no difference in *Col3a1* mRNA levels when the two cell states were compared in freshly isolated cells or after growth in GM (Figure 4G). Moreover, we monitored the expression of the *Timp1* mRNA, another transcript expressed by FAPs upon *in vitro* fibrogenic stimulation ^12^. SCA1-High-FAPs but not SCA1-Low-FAPs respond to the FM enhancing their expression of the *Timp1* mRNA (Figure 4F)

We conclude that both FAPs cell states are able to differentiate into myofibroblasts when exposed to a fibrogenic microenvironment, but they exhibit a differential expression of at least two genes coding an ECM component.

### SCA1-High-FAPs proliferate more extensively than SCA1-Low-FAPs

A coordinated modulation of FAP proliferation and apoptosis is a key requisite to achieve efficient muscle regeneration ^16^. Indeed, the deregulation of these processes observed in DMD leads to FAP accumulation and the ensuing deposition of ectopic tissues ^33^.

We investigated whether the observed FAP heterogeneity would also affect cell proliferation. Cells were sorted from *mdx* mice and the two cell states were plated at the same low density. Starting at the first time point (T0), we collected samples every 24 hours (Figure 5A) and we counted the number of nuclei and the percentage of Ki-67 expressing cells to identify proliferating cells. Already at 48 hours, the number of SCA1-High-FAPs nuclei in a field is significantly higher than in the SCA1-Low-FAPs (Figure 5B). However, the percentage of cells expressing Ki-67 is comparable in freshly purified cells and decreases more rapidly at later times in SCA1-High-FAPs as this cell state reaches confluence earlier and stops growing. In the exponential growth phase, the doubling time of SCA1-High-FAPs is about ten hours shorter (Supplementary Figure 7A and Figure 5D). Taken together these results suggest that *ex vivo* SCA1-High-FAPs have a shorter duplication time reaching at late time points a higher confluence and a lower number of Ki-67 positive cells.

**Figure 5.**
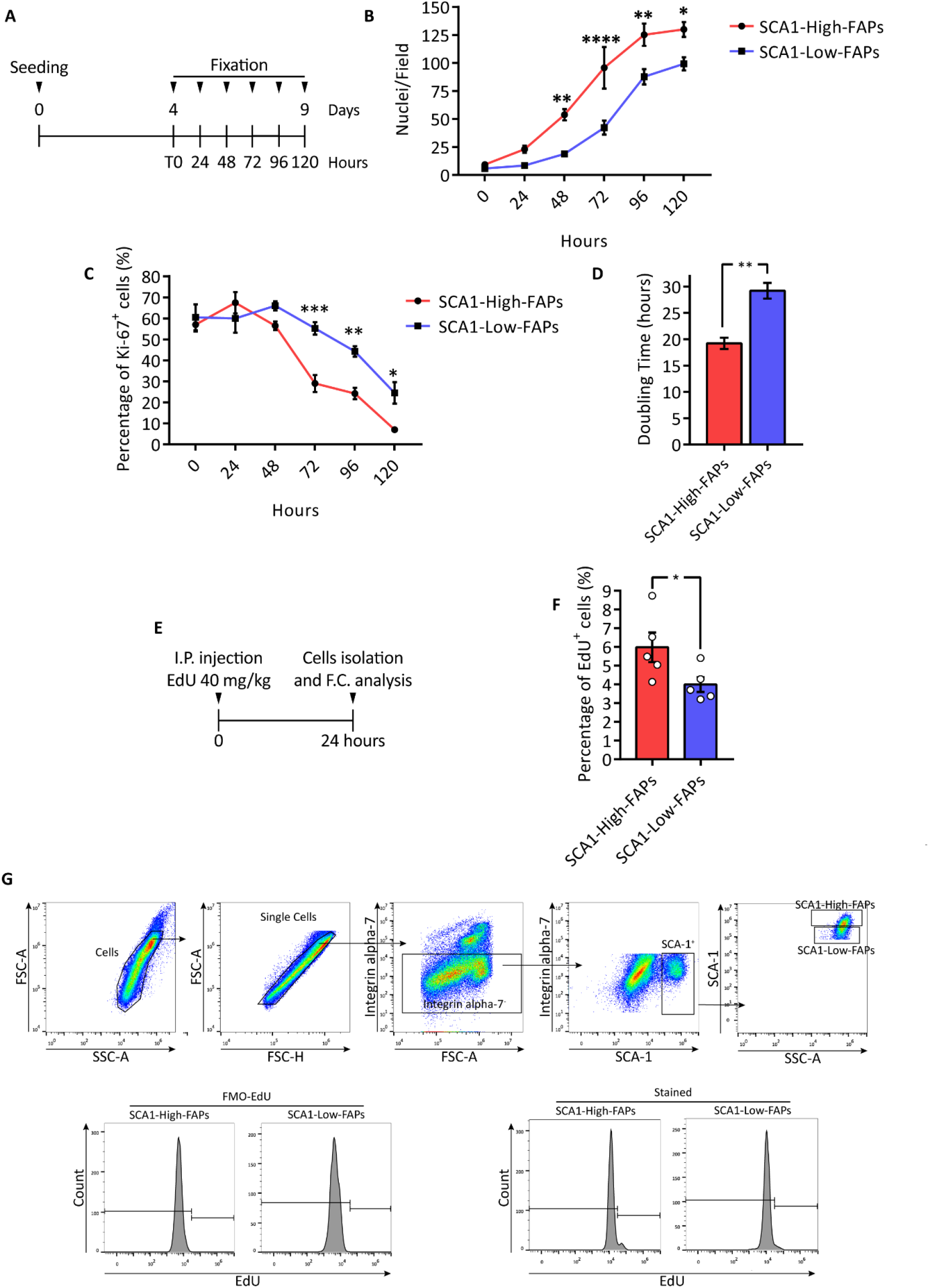
SCA1-High-FAPs have a higher proliferation rate. (A) Schematic representation of the experimental plan to study *mdx* FAP cell states proliferation rate *in vitro*. Collected samples were immunolabelled for Ki-67 and nuclei were counterstained with Hoechst 33342. n = 4. (B) Growth curve showing the number of nuclei per field. (C) Plot representing the percentage of Ki-67 positive nuclei per field. (D) Doubling time of FAP cell states calculated using the first three days of exponential growth. A Student t-test was applied to determine statistical significance. (E) Experimental plan applied to investigate cell states proliferation *in vivo*. EdU at the concentration of 40 mg/kg was injected into the peritoneum of *mdx* mice. After 24 hours mice were sacrificed and EdU incorporation was studied by flow cytometry. (F) Bar plot showing the percentage of FAP cell states in proliferating FAPs. n = 5. (G) Gating strategy to evaluate EdU incorporation. For the unstained sample and FMO controls see Supplementary Figure 7B-C. Data are presented as mean ± SEM. Statistical analysis was performed using Two-way ANOVA for the *in vitro* experiment and Student t-test for the *in vivo* experiment. * p ≤ 0.05, ** p ≤ 0.01, **** p ≤ 0.0001.

In order to lend support to an *in vivo* relevance of these observations, we injected 5-ethynyl-2’-deoxyuridine (EdU) into the peritoneum of *mdx* mice. After 24 hours we isolated mononuclear cells and we selected CD45^-^ CD31^-^ cells by magnetic activated cell sorting (MACS). Next, we identified FAP cell states in the Integrin alpha-7^-^ SCA-1^+^ compartment and monitored EdU incorporation using flow cytometry (Figure 5G; for the controls refer to Supplementary Figure 7B-C). A larger fraction of SCA1-High-FAPs have incorporated EdU in comparison with SCA1-Low-FAPs (6% versus 4%, p-value < 0.05) (Figure 5F). In conclusion, we observed a difference in the proliferation rate of FAP cell states *in vitro* which correlates with a higher fraction of SCA1-High-FAPs incorporating EdU *in vivo*.

### Single cell transcriptomic analysis of FAP cell states from wild type mice

Till now we have used the expression of a single antigen (SCA-1) to characterise two FAP cell states displaying distinct differentiation and proliferation characteristics. In these experiments cells were purified from muscles of *mdx* mice as in this model the two cell states are more evenly represented. However, heterogeneity in the expression of SCA-1 is observed both in wild type and *mdx* muscles (Figure 1). To obtain a more general picture of the observed heterogeneity by an independent experimental approach we made use of the Tabula Muris Senis dataset ^34^. The Tabula Muris Senis is a comprehensive compendium of single cell RNA sequencing datasets in 23 organs from wild type mice at different ages. We focused on the skeletal muscle and performed an unbiased automated clustering using markers from the Myo-REG resource (Supplementary Figure 8A) to identify the main skeletal muscle populations ^35^. We obtained 11 clusters (Figure 6A) with an expression profile characteristic of each population (Supplementary Figure 8B-L). Next, we carried out an additional clustering analysis on the FAP population and observed four clusters (Figure 6B) characterised by different levels of SCA-1 expression (Figure 6C). Cluster 0 (red), cluster 2 (blue) and cluster 3 (purple) express similar low level of SCA-1 and where collapsed into a single group, thus obtaining two populations expressing high and low SCA-1 levels (Figure 6D). SCA1-Low-FAPs are more numerous than SCA1-High-FAPs (Figure 6E), similarly to what was observed in young mice using the flow cytometry assay (Figure 1C). Moreover, their ratio only slightly increases in ageing.

**Figure 6.**
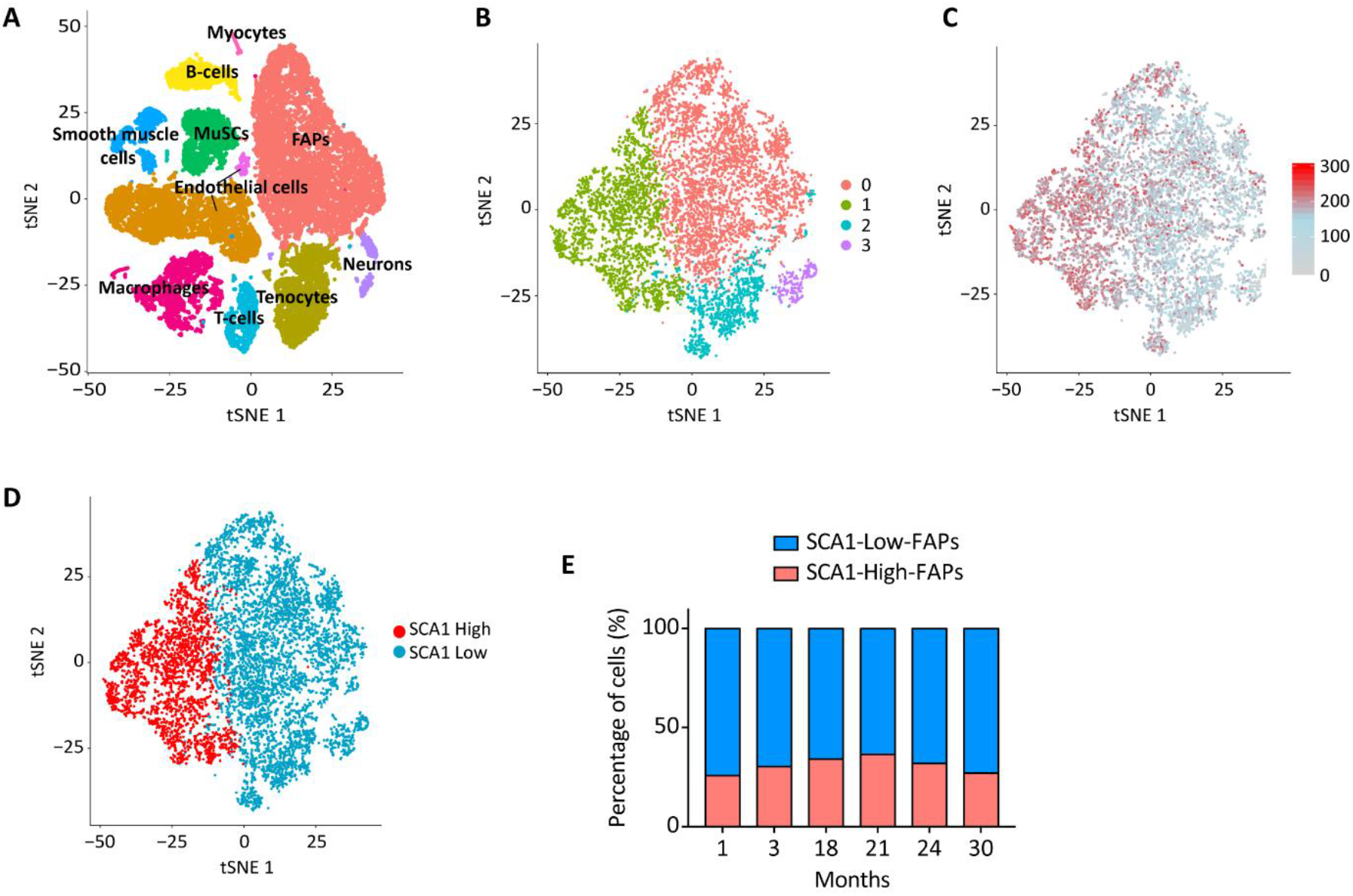
FAP cell states from wild type mice differ in their transcriptional profile. (A) viSNE map representing the different clusters identified as distinct cell populations according to the expression of specific biomarkers. Identified populations are: Myocytes, B-cells, FAPs, Smooth Muscle cells, MuSCs, Endothelial Cells, Macrophages, T-cells, Tenocytes and Neurons. (B) The FAPs population subset has been re-clustered, leading to 4 distinct clusters according to their gene expression profile. The four clusters (0, 1, 2, 3) were mapped onto the viSNE map of FAPs. (C) The expression of SCA-1 antigen has been mapped to each cell of the dataset. (D) viSNE map of FAPs in which clusters 0, 2, 3 are collapsed together to obtain two clusters expressing high level (SCA-1 High) and low level (SCA-1 Low) of SCA-1. (E) Stacked bar plot showing the percentage of the two population on the basis of SCA-1 expression levels across different ages.

We performed a Gene Ontology (GO) term enrichment analysis on differentially expressed genes which did not provide any robust clue regarding FAP cell states properties. However, the list of differentially expressed genes revealed that SCA1-High-FAPs express higher levels of *DPP4* mRNA which is a marker of mesenchymal progenitors residing in the reticular interstitium of inguinal white adipose tissue (iWAT) ^25^. In this tissue DPP4^+^ characterise progenitors able to differentiate into adipocytes suggesting an analogy between such progenitors and SCA1-High-FAPs. Next, we asked if the differential expression of the *COL1A1* gene observed in *mdx* FAP cell states (Figure 4) is conserved in the wild type counterpart. *COL1A1* mRNA is slightly downregulated in SCA1-High-FAPs (average log_2_ Fold Change = −0.341, adjusted p-value < 0.05), in accord with our findings in FAP cell states freshly isolated from *mdx* mice.

We conclude that single cell transcriptional profiling of wild type FAPs allow to distinguish two cell states on the basis of SCA-1 transcriptional levels. The analysis of genes that are differentially expressed in the two cell states provides hints about possible mechanisms underlying the observed phenotypic differences.

### Signals secreted from cells of the immune compartment control adipogenesis of FAP cell states in an age-dependent fashion

*In vivo* the FAP proliferation and differentiation traits are controlled by the muscle microenvironment ^13,14^. The prolonged and sustained chronic inflammation of *mdx* muscles is characterised by a subset of unconventional macrophages. It was shown that in an *in vivo* model recapitulating muscle aging in *mdx* mice, these macrophages fail to induce FAP apoptosis and promote their excessive growth ^33^. Others report that, in wild type mice, ageing determines a decrease in the number of FAPs associated with a reduced adipogenic potential ^36^.

To find out how ageing affects FAPs and FAP cell states in the *mdx* mouse we determined the number of FAPs in the TA of young and aged dystrophic mice by immunofluorescence (Figure 7A-B). We observed a reduction in the number of FAPs within old dystrophic muscles and this condition is followed by an increase of FAPs expressing PPAR-gamma in the tibialis anterior (TA) of aged *mdx* mice (Figure 7C). These results suggest that FAPs in the microenvironment of aged *mdx* mice are more committed to the adipogenic differentiation. To verify this hypothesis we set out to investigate the effect of the immune cell compartment of young and old *mdx* mice on the adipogenesis of FAP cell states. For this purpose, we isolated CD45^+^ cells (henceforth called leukocytes) from young (6-8 weeks) and old (60 weeks) *mdx* mice and after 24 hours of *in vitro* culture we harvested their supernatants. Next, we added each of these conditioned media (CM) to AIM and MM to test their effect on young FAP cell states differentiation (Figure 7D). Adipocytes were identified by ORO staining. The CM of leukocytes from young *mdx* mice significantly inhibits adipogenesis of SCA1-High-FAPs (~ 7-fold decrease) (Figure 7E-F). We observed a less important inhibition using the medium conditioned by leukocytes from old *mdx* mice (~3-fold decrease). In general, the differentiation potential of FAPs in both cell states is inhibited by the supernatants conditioned by leukocytes from young *mdx* mice and less so by supernatants of leukocytes from old *mdx* mice. While in these conditions the differentiation of SCA1-Low-FAPs is cut down to negligible levels, the differentiation of SCA1-High-FAPs still remains significant especially when exposed to conditioned media of leukocytes from old *mdx* mice. These observations suggest that the microenvironment of young *mdx* mice is more efficient in inhibiting FAP adipogenesis and probably sufficient to keep in check the differentiation of both FAP cell states. With aging, the negative control exerted by the CD45^+^ cell compartment becomes less effective, selectively releasing the adipogenic potential of SCA1-High-FAPs. These findings offer a possible mechanism to explain the appearance of IMAT in the TA of aged *mdx* mice as demonstrated by the increase of immunostaining against Perilipin (Figure 7G-H) and the increase of *Cebpb* mRNA in the whole TA (Figure 7I).

**Figure 7.**
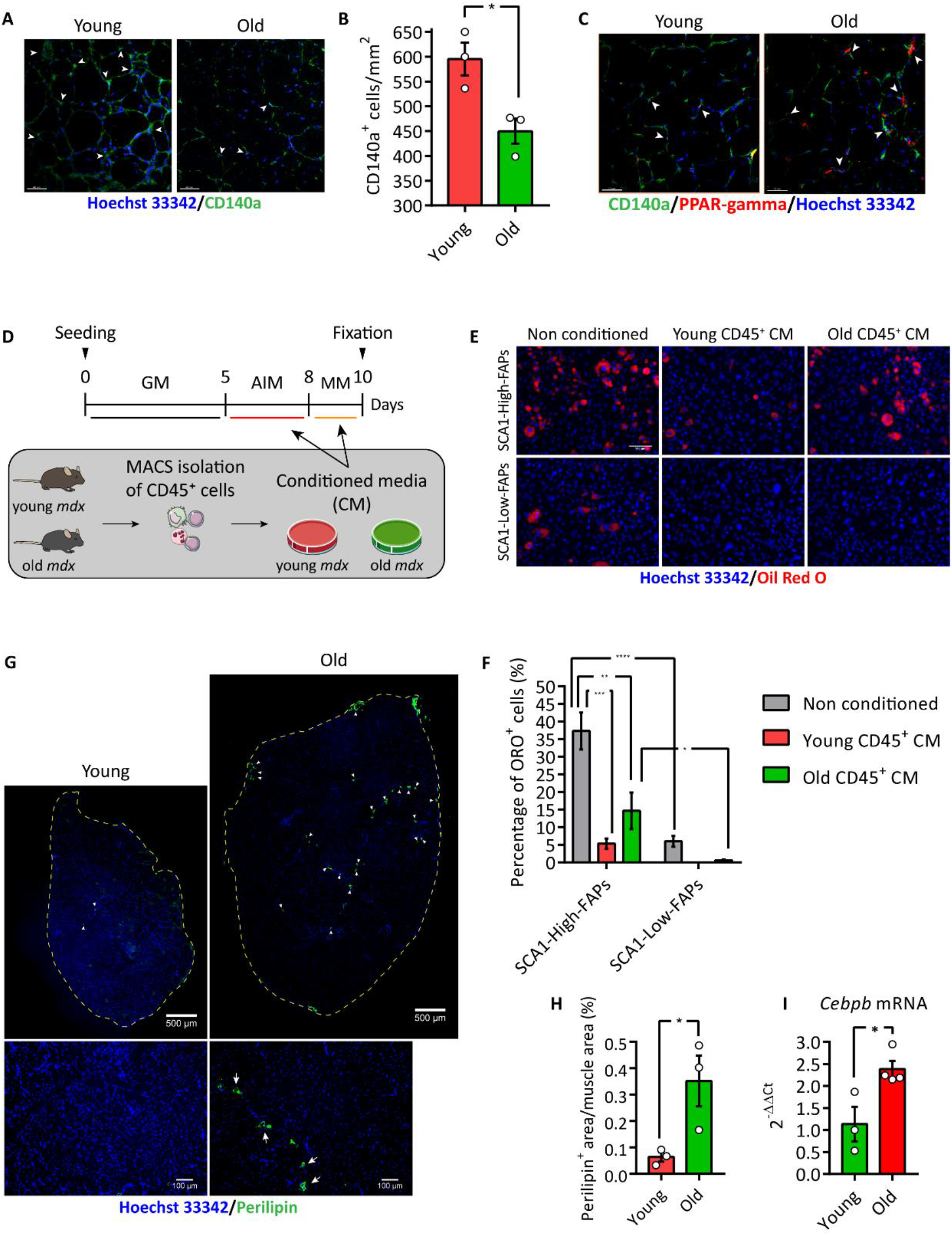
Immune system from old *mdx* mice releases adipogenic potential of SCA1-High-FAPs. (A) Representative images of the experiment whose results are reported in the bar plot in (B). FAPs were identified by an antibody against CD140a (green) and nuclei were counterstained with Hoechst 33342 (blue). Scale bar 40 μm. (B) Bar plot representing the number of FAPs (CD140a^+^ cells) per mm^2^ in a section of TA from young and old *mdx* mice. Statistical analysis was carried out by a Student t-test. n = 3. (C) Representative immunofluorescence of TA section of young and old mdx mice showing PPAR-gamma positive FAPs. FAPs were immunolabelled with an antibody against CD140a (green) and nuclei were counterstained with Hoechst 33342 (blue). In red cells expressing PPAR-gamma. Scale bar 40 μm. n = 3. (D) Schematic representation of the experiment in (E) and (F). Leukocytes (CD45^+^ cells) were isolated through MACS from young and old *mdx* mice. After 24 hours of *ex vivo* culture their conditioned media (CM) were harvested and used to induce adipogenesis of young FAP cell states from *mdx* mice in combination with AIM and MM. (E) Representative microphotographs of (D). (F) Bar plot showing the percentage of ORO positive cells. n = 3. Statistical significance was evaluated using One-way ANOVA. Lipid droplets were stained with ORO (red) and nuclei with Hoechst 33342 (blue). Scale bar 100 μm. (G) Immunofluorescence against perilipin on TA from young (left) and old (right) *mdx* mice. Representative images of reconstruction of the whole TA section. Perilipin is showed in green and nuclei in blue. Dashed lines highlight section borders. Scale bar 500 μm. Lower insets show images used to reconstruct the whole muscle. Scale bar 100 μm. Arrows indicate perilipin positive cells. (H) Bar plot presenting the percentage of Perilipin positive area in the whole TA section. Statistical significance was evaluated using a Student t-test. n = 3. (I) Bar plot showing the expression of *Cebpb* mRNA in the TA of young (n = 3) and old (n = 4) *mdx* mice assessed by Real-Time PCR using 2^-ΔΔCt^ comparative method. *Tubulin* mRNA was set as reference gene. Statistical significance was evaluated with a Student t-test. Data are presented as mean ± SEM. * p ≤ 0.05, ** p ≤ 0.01, *** p ≤ 0.001, **** p. ≤ 0.0001.

## Discussion

Non-genetic heterogeneity in clonal cell populations is suggested to underlie fundamental developmental and pathological processes ^7,21,37,38^. However, as a consequence of the dearth of appropriate technologies, this concept has not been widely addressed as many experimental approaches still rely on bulk analysis of cell populations. The emergence, over the past decade, of powerful single cell approaches has permitted to uncover a common hidden heterogeneity in biological systems ^39^. We have been interested in investigating whether a perturbation in the spread of a molecular profile may be associated to disease and whether conversely disease has an impact on population heterogeneity. To address this question, we applied a multiplex flow cytometry approach to study the expression of nine surface antigens in a suspension of primary cells from a solid tissue. The single cell expression profiles were used in a clustering approach to identify fibro/adipogenic progenitors (FAPs) in wild type and dystrophic mice. We confirmed the inherent heterogeneity of this cell population (Supplementary Figure 2) and by focusing on the expression of the SCA-1 antigen we identified two cell clusters that are present with different abundance in the muscles of wild type and *mdx* mouse models. Interestingly an unsupervised clustering of the single cell gene expression dataset of the Tabula Muris Senis Consortium also lead to similar conclusions identifying clusters of FAPs characterised by different expression levels of SCA-1.

Different kinds of cell to cell variability within a monoclonal cell population have been described using single cell technologies. In particular, the term micro-heterogeneity refers to a condition in which the distribution of the expression of a given trait, an mRNA or a protein, in a cell population is represented by a bell shaped curve whose standard deviation is larger than expected by random noise or experimental error ^6,40^. We identified such a condition in the expression distribution curve of SCA-1 in FAPs. SCA-1 micro-heterogeneity was already shown to modulate lineage choice in a Erythroid myeloid lymphoid (EML) cell line ^7,41^. This consideration encouraged us to further investigate the role of SCA-1 heterogeneity in fate decision of dystrophic FAPs. To this purpose, based on the results of our multiplex flow cytometry approach and prior knowledge ^7,8,21^, we resolved the continuity of the distribution of SCA-1 expression, a process defined decomposition ^40^, into two discrete groups of cells by a fluorescence activated sorting strategy based on SCA-1 expression levels.

Although efforts have been made, clear definitions and terminologies regarding sub-populations and cell states are still lacking. Here we use two *in vitro* criteria to distinguish these conditions: the nature of their heterogeneity and the ability to maintain their identity once they are separated and cultured *in vitro*. Here, for cell sub-populations, in a population displaying macro-heterogeneity, we mean two or more groups of cells in a bi-(multi-) modal distribution that maintain their identity when cultured *in vitro*. On the other hand, we define “cell states” group of cells deriving from a spread in a single peak distribution (micro-heterogeneity), but expressing different levels of a given molecular trait and failing to maintain their identity *in vitro*. By applying these criteria, the two groups of dystrophic FAPs were found to be morphologically identical cell states able to repopulate each other *in vitro*. Whether such definitions would have some functional relevance *in vivo* remains to be established.

The involvement of micro-heterogeneity in the decision of cell fate has been addressed in different biological contexts. In the pre-implantation embryo, blastomeres express variable level of CDX2 ^38^. In HeLa cells, the probability and timing of cell death in response to TRAIL are determined by the variance in protein levels of its signalling cascade ^37^. The role of stem cell heterogeneity during muscle regeneration was recognized only lately ^8^ and its contribution to FAP phenotype in Duchenne muscular dystrophy (DMD) has been highlighted even more recently ^17^. However, its role in the development of ectopic tissues in DMD is unexplored. Here we have reported our investigation on the impact of the observed FAP heterogeneity on differentiation and proliferation in FAPs isolated from the muscles of the *mdx* mouse, an animal model of DMD ^42^. FAPs are exposed *in vivo* to a complex and dynamic microenvironment which adds further complexity ^43^, hence initially we performed our investigation *in vitro* to take into account only their intrinsic SCA-1 heterogeneity and their response to a controlled microenvironment. We demonstrated that SCA-1 micro-heterogeneity affects the expression of PPAR-gamma in response to an adipogenic medium resulting in a higher propensity of SCA1-High-FAPs to fully differentiate into adipocytes. Moreover, despite FAP cell states showing a similar propensity to differentiate into myofibroblasts, SCA1-Low-FAPs do not increase their expression of *COL1A1* and *Timp1* mRNAs after fibrogenic stimulation. Interestingly, these cells express higher amount of *COL1A1* mRNA in both *mdx* and wild type mice. We surmise that the expression of *COL1A1* gene in SCA1-Low-FAPs do not respond to the fibrogenic medium probably because of an already high basal transcription. We also checked if SCA-1 micro-heterogeneity has an influence on FAP proliferation, a central process in muscle regeneration. The increase of approximately 10 hours in the doubling time of SCA1-Low-FAPs decrease their proliferation rate and lengthen the period in which cells are in proliferation. Overall, our results demonstrate that micro-heterogeneity in SCA-1 expression is involved in the fate decision of dystrophic FAPs in a controlled *in vitro* environment (Figure 8) and raise the possibility that this type of cell to cell variability may play a role in the differentiation and proliferation of mesenchymal-like cells in other pathological contexts.

**Figure 8.**
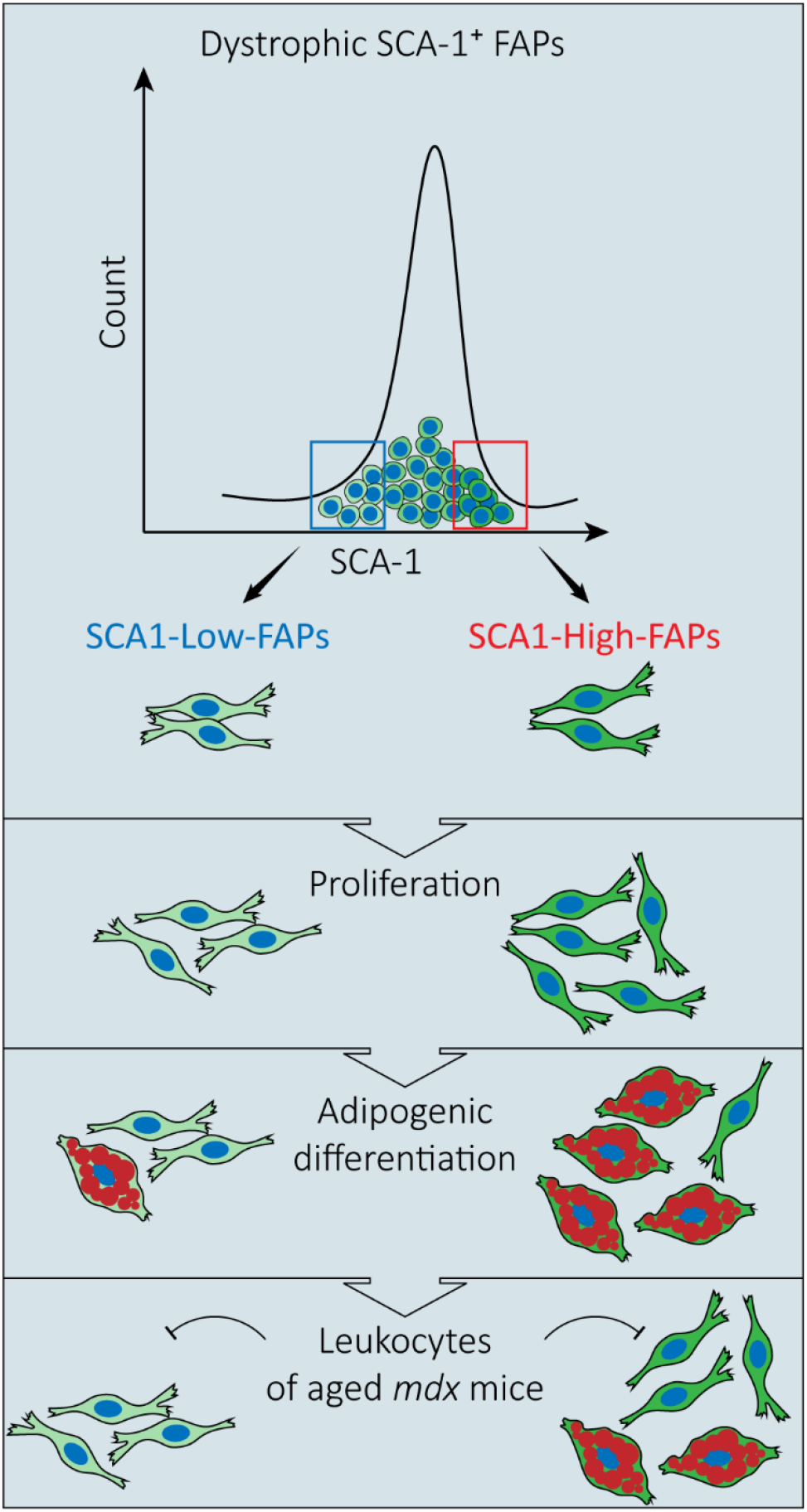
Schematic representation recapitulaning the role of SCA-1 micro-heterogeneity in FAP adipogenesis in dystrophic mice. Two FAP cell states expressing high and low levels of SCA-1 respectively were isolated from the hind limb muscles of mdx mice. The two cell states were tested for their differentiation and proliferation properties *in vitro*. SCA1-High-FAPs are more proliferating and more adipogenic cells in comparison with SCA1-Low-FAPs. Leukocytes from old dystrophic mice strongly inhibit adipogenesis in SCA1-Low-FAPs while are less effective on SCA1-High-FAPs.

We observe and characterise FAP heterogeneity in purified cells in the absence of stimuli from the muscle niche. This may lead to the conclusion that this micro-heterogeneity is a cell autonomous phenomenon that is accentuated by the mutation in the dystrophin gene. It is however unlikely that such a mutation alters the expression of the SCA-1 antigen and promotes a diversity in the adipogenic potential of different FAP cells.

The challenge is now to explore the physiological relevance of these observations *in vivo* where additional complexity is added by the interaction of micro-heterogeneity with a dynamic microenvironment. To provide some evidence in this respect, we analysed the proliferation of FAPs by looking at the incorporation of EdU after injection into the peritoneum of *mdx* mice. The results confirmed that *in vivo*, as in isolated cells, FAPs expressing different level of SCA-1 proliferates to a different extent.

Our results show an increase of Perilipin and *Cebpb* mRNA in old *mdx* mice muscles associated to a decreased number of FAPs. These FAPs are also more committed to adipogenic differentiation as demonstrated by the increase of PPAR-gamma positive FAPs. We considered the possibility that the microenvironment under the stress caused by the muscle chronic damage could affect FAP micro-heterogeneity and differentially promote or inhibit the adipogenic potentials of FAP expressing high and low levels of SCA-1. To this end we repeated the differentiation experiment using adipogenic media conditioned by leukocytes (CD45^+^ cells) from young and old *mdx* mice. In this condition, the two FAP cell states respond differently to the stimuli in the leukocyte supernatants. Adipogenesis of both cell states is inhibited by leukocyte secreted factor(s). The supernatant of leukocytes from young *mdx* mice is more efficient in restraining adipogenesis of both FAP cell states consistent with the observation that no fat infiltrations are observed in young *mdx* mice. With aging, however, leukocytes become less efficient in inhibiting adipogenesis of SCA1-High-FAPs (Figure 8). These conditions are still far from closely matching the *in vivo* microenvironment of young and old dystrophic muscles, as for instance the regulatory mechanisms provided by the muscle fibres are not included ^18^. However, these observations, suggesting a different age dependent response of FAP cell states to the anti-adipogenic signals of the immune compartment may explain the increase of IMAT in muscles of old dystrophic mice.

In conclusion, we surmise that micro-heterogeneity in SCA-1 expression plays a role during the decision-making of *mdx* FAPs *in vitro* and it is affected by the environment of dystrophic muscles.

## Materials and Methods

### Mouse strains

C57BL/6J (wild type) and C57BL10ScSn-Dmd^mdx^/J (*mdx*) mice were purchased from the Jackson Laboratories. We employed young (1.5 months old) and old (15 months old) mice. Whenever possible, we balanced gender of the mice in each experiment. Animals were bred and maintained according to the standard procedures of the animal facility. The experimental procedures were conducted according to the rules of good animal experimentation I.A.C.U.C. n 432 of March 12 2006 and under the approval released from the Italian Ministry of Health.

### Isolation of primary cells

Wild type and *mdx* mice were sacrificed through cervical dislocation and were washed with 70% ethanol. After an incision through the skin, hind limbs were excised and placed into cold Hank’s Balanced Salt Solution without Calcium and Magnesium (HBSS, Biowest L0605-500) supplemented with 0.2% bovine serum albumin (BSA, Applichem PanReac A1391) and 100 U/ml penicillin, 100 mg/ml streptomycin (Gibco 15140122). Hind limb muscles were removed from bones under sterile hood and mechanically minced using a scalpel. Minced tissue was washed with HBSS and centrifuged at 700 x g for 10 minutes at 4 °C. Pelleted tissue was weighted and resuspended in the enzymatic digestion mix composed by 2,4 U/ml dispase II (4 ml/g of muscles) (Roche 04942078001) dissolved in Dulbecco’s phosphate buffered saline (D-PBS) with Calcium and Magnesium (Biowest L0625-500), 0,01 mg/ml DNase I (Roche 04716728001) and 2 μg/ml collagenase A (Roche 10103586001). Tissue preparations were incubated at 37 °C in a water bath (in gently shaking and not in immersion) for 1 hour vortexing every 30 minutes. Digestion was stopped by the addition of HBSS and samples were centrifuged at 700 x g for 10 minutes at 4 °C. Pellets of cells were resuspended in 10 ml of HBSS and filtered through a 100 μm cell strainer (Falcon 352360). Cell suspensions were centrifuged at 700 x g for 10 minutes at 4 °C, pellets were resuspended in 10 ml of HBSS and filtered through a 70 μm cell strainer (Falcon 352350). After a further centrifuge red blood cells were lysed in 1 ml of RBC lysis buffer (Santa Cruz Biotechnology sc-296258) in ice for 2.5 minutes. Lysis was stopped adding HBSS and cell suspensions were filtered through a 40 μm cell strainer (Falcon 352340). Cell strainers were always washed before and after the use with 5 ml of HBSS. Cell suspensions were centrifuged at 700 x g for 10 minutes at 4 °C and resuspended in 500 μl Magnetic Beads Buffer (MBB) composed by D-PBS without Calcium and Magnesium, 0.5% BSA and 2 mM Ethylenediaminetetraacetic acid (EDTA). Cells were filtered through a 30 μm cell strainer (Miltenyi 130-041-407) which was washed 3 times with MBB. Mononuclear cells in this suspension was counted and centrifuged at 700 x g for 10 minutes at 4 °C. The isolation protocol proceeds with the magnetic activated cell sorting (MACS) of the CD45^-^ CD31^-^ cells. Pellets were resuspended in MBB and incubated with a microbead conjugated antibody against CD45 (Miltenyi 130-052-301) according to manufacturer’s instructions. After 15 minutes, cells were washed with 2 ml of MBB and centrifuged. Pellets were resuspended in 500 μl of MBB and cells were separated with MS columns (Miltenyi 130-042-201) according to manufacturer’s instructions to collect CD45^-^ cells. Protocol proceeds with the selection of CD31^-^ cells applying the same procedures described for CD45^-^ cells. Next, FAPs and muscle satellite cells (MuSCs) were isolated through fluorescent activated cell sorting (FACS). CD45^-^ CD31^-^ cell suspensions were incubated with the following primary antibodies for 30 minutes at 4 °C in D-PBS without Calcium and Magnesium, 2% BSA and 2 mM EDTA at 1 x 10^6^ cells per 100 μl: anti-ITGA7 APC 1:500 (Invitrogen MA5-23555) and anti-SCA-1 FITC 1:50 (BD Pharmingen 557405). Gates were strictly designed using proper fluorescence minus one (FMO) controls for each antibody. Debris and cell clusters were excluded using side scatter (SSC) and forward scatter (FSC) as showed in supplementary figures. Morevoer, we applied two criteria to enhance the reproducibility of the sorting strategy of FAP cell states: FAP cell states must represent at least 80% of the total FAPs (this criteria also increase the yield of the isolation); the mean intensity of SCA-1 must differ of at least 3-fold between the two cell states. FAPs and MuSCs were sorted directly in their growth media as ITGA7^-^ SCA-1^+^ and ITGA7^+^ cells respectively. Purity of sorted cells was assessed immediately after sorting.

Sorting was performed on FACSAria™ III (Becton Dickinson) cell sorter equipped with 5 lasers (18 parameters). Analysis were carried out using FlowJo™ Software.

### EdU administration

5-ethynyl-2’-deoxyuridine (EdU) (Invitrogen, C10418) was dissolved with 2 ml sterile PBS at the concentration of 5 mg/ml (20 mM). Mice were weighted and injected intraperitoneally with 40 mg/kg EdU ^33^. After 24 hours, CD45^-^ CD31^-^ were isolated by MACS and staining were performed using Click-iT− EdU Pacific Blue™ Flow Cytometry Assay Kit (Invitrogen, C10418). Briefly, 1 x 10^6^ CD45^-^ CD31^-^ were incubated with anti-ITGA7 APC 1:500 (Invitrogen MA5-23555) and anti-SCA-1 FITC 1:50 (BD Pharmingen 557405) antibodies for 30 minutes in ice. The subsequent fixation, permeabilization and Click-iT− reaction were performed as indicated by manufacturer’s instructions. Unstained cells and FMO controls were prepared. Samples were resuspended at the concentration of 1x 10^6^ cells per ml in PBS 1% BSA and analysed as described in the flow cytometry section.

### Flow cytometry

Cell suspensions were incubated with primary antibodies in ice for 30 minutes in PBS containing 2 mM EDTA and 2% BSA at the concentration of 1 x 10^6^ cells/ml. We used the following antibodies and dilutions: anti-ITGA7 APC 1:500 (Invitrogen MA5-23555), anti-SCA-1 FITC 1:50 (BD Pharmingen 557405) and anti-CD140a APC 1:50 (eBioscience 17-1401). To stop the incubation cell suspensions were diluted with 2 ml of PBS 2 mM EDTA 2% BSA and centrifuged at 4 °C for 10 minutes at 700 x g. Pellets were resuspended in PBS 2 mM EDTA 2% BSA at the concentration of 1 x 10^6^ cells/ml. Unstained samples and suitable FMO-controls were prepared when necessary. Approximately 10,000 events per samples were acquired by CytoFLEX S (Beckman Coulter) equipped with three lasers (488 nm, 405 nm and 638 nm) and 13 detectors. Quality control of the cytometer was assessed daily using CytoFLEX Daily QC Fluorospheres (Beckman Coulter B53230). Data were collected by CytExpert (Beckman Coulter) software. If needed, a compensation matrix was calculated using VersaComp Antibody Capture Kit (Beckman Coulter B22804) according to manufacturer’s instructions. FCS files were analysed using FlowJo™ Software.

### Multiplex flow cytometry assay and analysis

Total mononuclear cells were isolated from wild type and *mdx* mice as described. 1 x 10^6^ cells were stained in PBS containing 2 mM EDTA and 2% BSA at the concentration of 1 x 10^6^ cells/ml in ice for 30 minutes with the following antibodies: anti-CD140b Super Bright 436, 1 μg/100 μl (eBioscience 62-1402-82); anti-CD140a Super Bright 600, 0.5 μg/100 μl (eBioscience 63-1401-82); anti-CD31 Super Bright 645, 0.25 μg/100 μl (eBioscience 64-0311-82); anti-CD34 FITC, 1 μg/100 μl (eBioscience 11-0341-82); anti-CD146 PE, 0.125 μg/100 μl (eBioscience 12-1469-42; anti-SCA-1 PE-eFluor 610, 0.5 μg/100 μl (eBioscience 61-5981-82); anti-ITGA7 APC, 10 μl/1 x 10^6^ cells (Invitrogen MA5-23555); anti-CD45 Alexa Fluor 700, 0.5 μg/100 μl (eBioscience 56-0451-82); anti-CD90.2 APC-eFluor 780, 0.125 μg/100 μl (eBioscience 47-0902-82). 2 ml of PBS 2 mM EDTA 2% BSA was added to stop the incubation, cell suspensions was centrifuged and cell pellets were suspended in in PBS containing 2 mM EDTA and 2% BSA. At least 300,000 events were acquired by CytoFLEX S (Beckman Coulter) and collected CytExpert (Beckman Coulter) software. Wild type and *mdx* mice samples were acquired at different days therefore we created two different compensation matrixes. Next FCS files of these two datasets were simultaneously uploaded on Cytobank platform to perform multi-dimensional analysis. Cells were identified by SSC-A and FCS-A and single cells by FCS-A and FSC-H applying ‘tailored’ gates for each sample.

Dimensional reduction was carried out applying the tSNE ^44^ algorithm as followed: 50,000 events per sample; 2,000 iterations; perplexity = 30; theta = 0.5. All 9 channels were selected applying compensation matrixes.

To identify main muscle populations firstly we ran the FlowSOM algorithm with the following settings: all markers were selected; 50,000 events per sample; 100 clusters; 25 metaclusters; 10 iterations with scale normalisation. A hierarchical consensus clustering method was used to obtain the metaclusters. Next, metaclusters with similar expression profiles were merged together using the ‘automated cluster gates’ function.

FAP heterogeneity were studied applying the FlowSOM algorithm as followed: 600 events per samples; selected markers were specified in the results section; 100 clusters; 4 metaclusters; 10 iterations with scale normalisation. A hierarchical clustering method was used to obtain the metaclusters. We merged the two metaclusters expressing higher levels of SCA-1 and the two metaclusters expressing lower levels of SCA-1 obtaining two groups of FAPs. Self-Organising Maps (SOM) were uploaded in Cytoscape ^45^ and graph layouts were modified.

### Mass cytometry

Isolation of FAPs by MACS technology from *mdx* mice was performed according to the protocol described in Cerquone Perpetuini et al., 2020.

FAPs were incubated with Cell-ID-Cisplatin-194Pt (Fluidigm) at the concentration of 1 μM to stain dead cells. Next, samples from three independent biological replicates were barcoded with the a combination of palladium isotopes of the Cell-ID 20-Plex Pd Barcoding kit (Fluidigm) at the concentration of 5 μM. Staining was quenched by adding Maxpar cell staining buffer (Fluidigm). Barcoded samples were mixed and stained with lanthanides-conjugated antibodies according to the manufacturer’s instructions. The full list of antibodies purchased from Fluidigm is: anti-CD45 89Y (3089005B); anti-CD146 141Pr (3155006B); anti-cleaved CASP-3 142Nd (3142004A); anti-CD34 144Nd (3143009B); anti-phospho-EGFR 146Nd (3146007A); anti-CD140a 148Nd (3148018B); anti-pRB 150Nd (3150013A); anti-CD140b 151Eu (3151017B); anti-Vimentin 154Sm (3154014A); anti-CD90.2 156Gd (3156006B); anti-phospho-STAT3 158Gd (3158005A); anti-CXCR-4 159Tb (3159030B); anti-Integrin alpha-7 161Dy (67-0010-05); anti-SCA-1 164Dy (3164005B); anti-phospho-CREB-1 176Yb (3176005A); DNA1 191Ir and DNA2 193Ir (201192A); Cisplatin 195Pt (201194); anti-CD31 165Ho (3165013B); anti-Actin 175Lu (3175026A); anti-c-kit 166Er (3166004B). Next, the suspension was incubated with Cell-ID Intercalator-Ir (Fluidigm) at the concentration of 125 nM for 1 hour in Maxpar fix and Perm Buffer (Fluidigm). Finally, cells were filtered with 30 μm cell strainer.

Samples were acquired using CyTOF2 tuned and calibrated according to manufacturer’s instructions. Rate of acquisition was < 400 events per second. Dataset was converted in .fcs files using Debarcoding software (Fluidigm) and normalised.

### Analysis of mass cytometry data

Debarcoded .fcs files were uploaded to Cytobank platform. Cells were identified using DNA1 and DNA2 content identified by Ir intercalator, while single cells were identified using the event length parameter and DNA1 (or DNA2). Next, dead cells were gated out using Pt channel.

For the experiment with FAPs isolated by MACS and induced to differentiate into adipocyte (Figure 3H-I), dimensional reduction was carried out applying the tSNE algorithm as followed: 35,000 events per sample; 1,000 iterations; perplexity = 30; theta = 0.5. All parameters were selected as clustering channels. To identify FAP cell states we ran the FlowSOM algorithm with the following settings: SCA-1 and phospho-CREB-1 were selected as clustering channels; all events were selected; 100 clusters; 3 metaclusters; 10 iterations. A k-Means clustering method was used to obtain the metaclusters. Next, metaclusters with similar expression profiles were merged together using the ‘automated cluster gates’ function and mapped onto viSNE maps to facilitate results interpretation.

### Cell Culture

Freshly isolated *mdx* FAPs cell states were seeded on 96 well plates (Falcon 353072) in growth medium (GM) at the density of 15,000 cell/cm^2^ for differentiation assays and at the density of 6,200 cells/cm^2^ for the proliferation assay. GM is composed by Dulbecco’s Modified Eagle Medium, High Glucose, GlutaMAX™ (Gibco 61965-026) supplemented with 20% fetal bovine serum (FBS) (Euroclone, ECS0180L), 100 U/ml penicillin and 100 mg/ml streptomycin (Gibco 15140122), 1 mM sodium pyruvate (Sigma-Aldrich S8636) and 10 mM 4-(2-hydroxyethyl)-1-piperazineethanesulfonic acid (HEPES) (Sigma H0887).

To induce adipogenic differentiation cells were exposed to the adipogenic induction medium (AIM) consisting of GM complemented with 1 μg/ml insulin (Sigma-Aldrich I9278), 1 μM dexamethasone (Sigma D4902) and 0.5 mM of 3-isobutyl-1-methylxanthine (IBMX) (Sigma, I5879). After three days, cells were exposed to the maintenance medium (MM) composed by GM and 1 μg/ml insulin for additional two days.

To induce fibrogenic differentiation cells were cultured in a medium composed by Dulbecco’s Modified Eagle Medium, High Glucose, GlutaMAX™, 100 U/ml penicillin and 100 mg/ml streptomycin, 1 mM sodium pyruvate and 10 mM HEPES, 5% horse serum (HS) (Euroclone ECS0090D) and 1 ng/ml TGF-beta-1 (PreproTech 100-21). To study the expression of collagen genes, cells were plated in 24 well plates at the concentration of 15,000 cells/cm^2^.

For *in vitro* expansion, FAPs cell states were cultured at the concentration of 3,400 cells/cm^2^ for 9 days in Cytogrow (Resnova TGM-9001-B) medium. Afterwards, cell monolayers were washed twice in sterile D-PBS and cells were detached with Trypsin 0.5 g/L EDTA 0.2 g/L (Lonza, # BE17-161E) for 5 minutes. Next, cells were processed for flow cytometry analysis.

Muscle satellite cells (MuSCs) were cultured in a growth medium composed by Dulbecco’s Modified Eagle Medium, High Glucose, GlutaMAX™, 100 U/ml penicillin and 100 mg/ml streptomycin, 1 mM sodium pyruvate, 10 mM HEPES, 20% FBS, 10% Horse serum (Euroclone ECS0090D), 2% Chicken embryo extract (Seralab CE-650-J).

CD45^+^ leukocytes were isolated from *mdx* mice with MACS technology. They were cultured for 24 hours in DMEM Dulbecco’s Modified Eagle Medium, High Glucose, GlutaMAX™, 100 U/ml penicillin and 100 mg/ml streptomycin with 0.2% BSA ^46^ at 375,000 cells/ml. Next, media were harvested, centrifuged at 700 x g for 10 minutes at 4 °C and finally supernatant were filtered to remove cell debris. Conditioned media from different biological replicates were mixed just before the treatment. One volume of the resulting conditioned medium was mixed with one volume of 2X AIM to induce adipogenesis. The same procedure was applied to prepare MM.

### Immunofluorescence

Indirect immunolabelling was performed as followed. Cells were fixed with 2% paraformaldehyde (Santa Cruz sc-281692) for 20 minutes at room temperature (RT). Paraformaldehyde was removed by inversion and cells were washed three times in PBS and one time in PBS 0.5% Triton X-100. Permeabilization was performed incubating cells for 5 minutes with 0.5% Triton X-100 (Sigma T9284) in PBS. Permeabilization solution was removed by inversion and cells were washed with 0.1% Triton X-100 in PBS. Unspecific sticky sites were blocked with a blocking solution consisting of 10% FBS 0.1% Triton X-100 in PBS for 30 minutes at RT. Next, samples were incubated with the following primary antibodies for 1 hour at RT: rabbit anti-PPAR-gamma 1:200 (Cell Signaling 2443S), mouse anti-SMA 1:300 (Sigma A5228) and rabbit Ki-67 1:400 (Cell Signaling 9129). For CD140a immunolabelling, cells were incubated over night at 4 °C with the anti-CD140a (R&D Systems AF1062) diluted 1:80 in blocking solution. Then samples were washed three times with 0.1% Triton X-100 in PBS for 5 minutes and were incubated 30 minutes at RT with the following secondary antibody diluted 1:200: goat anti-rabbit Alexa Fluor 488 (Southern Biotech 4050-30), goat anti-mouse Alexa Fluor 488 (Southern Biotech 1030-30) donkey anti-goat Alexa Fluor 488 (Southern Biotech 3425-30). Samples were washed three times with PBS 0.1% Triton X-100 for 5 minutes and nuclei were counterstained with 2 μg/μl Hoechst 33342 (Invitrogen H3570) in PBS 0.1% Triton X-100 for 5 minutes at RT. Finally, cells were washed three times in PBS for 5 minutes and samples were stored at 4 °C in PBS containing 0.02% of sodium azide.

For immunolabelling on muscle section, tibialis anteriors (TA) were embedded in optical cutting temperature (OCT) compound (Bio-optica 05-9801) and snap frozen in liquid nitrogen. Embedded tissues were stored at −80 °C. TA were sectioned using a cryostat (Leica) to produce transverse sections of 8 μm thickness. Sections were stored at −80 °C. Sections were fixed in 4% PFA for 10 minutes at RT and washed three times with PBS for 5 minutes. Next, permeabilization was performed incubating sections with 0.3% Triton X-100 in PBS for 30 minutes at RT. Unspecific sticky sites were blocked by a blocking solution composed of 0.1% Triton X-100 1% glycine (SERVA 23390.02) 10% goat serum (Euroclone ECS0200D) in PBS for 1 hour at RT. Anti-perilipin antibody (Cell signalling 3470) was diluted 1:100 in blocking solution and incubated with sections over night at 4 °C. Sections were washed twice in 0.2% Triton X-100 1% BSA in PBS (washing solution) for 15 minutes and 5 minutes at RT. Incubation with goat anti-rabbit Alexa Fluor 488 1:200 (Southern Biotech 4050-30) diluted in blocking solution was performed at RT for 30 minutes. At the end of incubation, sections were washed 15 minutes in washing solution and 5 minutes in PBS. Finally, nuclei were stained with 2 μg/μl Hoechst 33342 (Invitrogen H3570) in PBS for 10 minutes at RT. Sections were washed twice in PBS for 5 minutes. Finally, slides were covered with coverslips using Aqua-Poly/Mount (Tebu-bio 18606-20) and stored at 4 °C.

For the on-section detection of FAPs, TA section were fixed in 2% PFA and washed three times with PBS. Permeabilization was performed incubating sections with 0.3% Triton X-100 in PBS for 30 minutes. This step was directly followed by an incubation with Protein Block Serum Free (Dako X0909) for 30 minutes to block the sticky sites. After a wash in PBS for 5 minutes, sections were incubated over night at 4 °C with the following antibodies diluted in PBS 2.5% BSA: goat anti-CD140a 1:80 (R&D Systems AF1062) and rabbit anti-PPAR-gamma 1:200 (Cell Signaling 2443S). The following day section were washed three times for 10 minutes with PBS 0.1% Triton X-100. Next, samples were incubated for 30 minutes with the following secondary antibodies diluted in PBS 2.5% BSA: donkey anti-goat Alexa Fluor 488 1:200 (Southern Biotech 3425-30) and donkey anti-rabbit Alexa Fluor 555 1:200 (Thermo fisher scientific A32794). After two wash for 5 minutes each with PBS 0.1% Triton X-100 and three wash in PBS, section were incubated with 2 μg/μl Hoechst 33342 (Invitrogen H3570) in PBS for 10 minutes at RT. Sections were washed three times in PBS for 5 minutes. Finally, slides were covered with coverslips using Aqua-Poly/Mount (Tebu-bio 18606-20) and stored at 4 °C.

### Oil red O staining

Oil red O (ORO) (Sigma-Aldrich O0625) was dissolved in isopropanol at the concentration of 3.5 mg/ml. This stock was diluted to a working solution composed of 3 volumes of ORO and 2 volumes of distilled water. This ORO solution was filtered two times just before the use. Fixed cells were washed three times in PBS for 5 minutes and was incubated with ORO for 10 minutes at RT. ORO was removed by inversion and cells were washed three times in PBS for 5 minutes. Nuclei were counterstained with 2 μg/μl Hoechst 33342 in PBS 0.1% Triton X-100 for 5 minutes at RT. Finally, cells were washed three times in PBS for 5 minutes and were stored at 4 °C in PBS containing 0.02% of sodium azide.

During immunofluorescence, ORO staining was performed after incubation with secondary antibodies. After three washes in PBS protocol continues with the Hoechst 33342 incubation as described in immunofluorescence section.

### CFSE staining and analysis of cell morphology

CarboxyFluorescein Succinimidyl Ester (CFSE) (Abcam ab113853) staining was carried out according to manufacturer’s instructions. Culture medium was removed and then 10 μM of CFSE in PBS was overlaid onto cells. After 15 minutes of incubation at 37 °C, staining was quenched adding an equal volume of GM and allow to sit for 5 minutes at 37 °C. Next medium was removed and cells were washed one time in PBS and fixed with 2% paraformaldehyde. Samples were washed three times in PBS for 5 minutes. Nuclei were counterstained with 2 μg/μl Hoechst 33342 in PBS 0.1% Triton X-100 for 5 minutes at RT. After three wash in PBS samples were at 4 °C in PBS containing 0.02% of sodium azide.

Images were manually acquired. Images were analysed using Fiji. Background signal was removed using the subtract background command and an image specific threshold was applied to obtain a binary image. Then we run the following series of commands to define cell borders: median filter, erode and fill holes. Finally, we measured area occupied, roundness and aspect ratio. We analysed at least 50 cells ^23^ from a total of three independent biological replicates

### Acquisition of images and analysis

Images were acquired with a LEICA microscope (DMI6000B) using LAS X software. For experiments using *ex vivo* cell culture, a matrix of 25 non-overlapping field per well was acquired. Experiments were performed in technical duplicate. ORO positive cells, Ki-67 positive nuclei and PPAR-gamma positive nuclei were manually counted using Fiji and were divided by the number of nuclei per field. SMA-positive area per cell was assessed using a dedicated pipeline of CellProfiler ^47^. Area expressed in pixel was converted in square micron and normalized over the total number of nuclei in each field.

Microphotographs of TA sections immunolabelled for perilipin were acquired using a 10X objective. At least three section for each mouse were acquired. These overlapped images were used to reconstruct the whole TA section through the Grid/collection stitching algorithm available in Fiji. To identify perilipin positive area we applied a threshold (0-100) and we divided this area by the whole section area.

Microphotographs of TA section labelled for CD140a were acquired with Leica microscope for the quantification and representative images were acquired using the confocal microscope Olympus IX-81 with a 40X objective.

Images of CD140a and PPAR-gamma positive FAPs on TA section were acquired using the confocal microscope Olympus IX-81 with a 40X objective.

### RNA extraction, retrotranscription and Real-Time PCR

Cell monolayers were washed in PBS and lysed in TRIzol™ Reagent (Invitrogen 15596-018). For extraction from tissue, TA were homogenised in liquid nitrogen using a pestle and then resuspended in TRIzol™. Isolation of total RNA were performed according to manufacturer’s instruction. RNA precipitation was performed at −20 °C overnight with glycogen. Total RNA was resuspended in nuclease free water, concentration and 260 nm/280 nm ratio were determined by Nanodrop Lite Spectrophotometer (Thermo Fisher Scientific). Samples were diluted to the concentration of 50 ng/μl and stored at −80 °C. cDNA was generated by PrimeScript RT Reagent Kit (Takara RR037A). Real-time PCR was performed on 15 ng of cDNA using SYBR Premix Ex Taq (Tli RNaseH Plus) (Takara RR420W) in a reaction volume of 20 μl. Tubulin was used as housekeeping gene. We used the following primers: *Tubulin* forward 5’-AAGCAGCAACCATGCGTGA-3’; *Tubulin* reverse 5’-CCTCCCCCAATGGTCTTGTC-3’; *Col1a1* forward 5’-GCAACAGTCGCTTCACCTACA-3’; *Col1a1* reverse 5’-CAATGTCCAAGGGAGCCACAT-3’; *Col3a1* forward 5’-CAACGGTCATACTCATTC-3’; *Col3a1* reverse 5’-TATAGTCTTCAGGTCTCAG-3’; *Timp1* forward 5’-AGCCTCTGTGGATATGCCC-3’; *Timp1* reverse 5’-TCAGAGTACGCCAGGGAACC-3’; *Cebpb* forward 5’-CAAGCTGAGCGACGAGTACA-3’; *Cebpb* reverse 5’-GACAGCTGCTCCACCTTCTT-3’. Comparative expression was performed using the 2^-ΔΔCt^ method ^48^ where ΔΔC_t_ is: (C_t, target_ − C_t, tubulin_) − (C_t, mean target_ − C_t, mean tubulin_).

### Single-cell RNAseq data analysis

Briefly, single cell RNAseq data of hind limb muscle have been retrieved from Tabula Muris Senis project and re-processed with the Seurat R package ^49^. After log normalization, we used the 2,000 most variable features to proceed to downstream analyses. Data have been scaled and the first 10 principal components used to run the t-distributed stochastic neighbour embedding (tSNE) algorithm. Whenever possible, biomarkers have been retrieved from Myo-REG database to assign cell line ontology identifications to each cluster, on the basis of the expression level of each biomarker for each cluster, according to their distribution (Supplementary Figure 8).

Once cell IDs have been assigned, FAPs population subset has been further analysed for the expression of SCA-1 and downstream analyses.

### Gene set enrichment analysis

Differentially expressed genes were identified by comparing the mRNA expression profile of the two FAPs clusters. Only genes whose expression difference had an adjusted p-value < 0.05 were considered as sub-population biomarkers. EnrichR ^50^ has been used to retrieve the enriched GO Molecular Function terms for each population with p-value < 0.05.

### Statistical analysis

Experiments were performed with primary cells isolated from at least three different mice (here defined biological replicates). Number of biological replicates were specified in the figure legends. Statistical analysis was performed by a two tailed Student t-test between two groups, while one-way ANOVA or two-way ANOVA were applied for comparison among more than three groups. Results were presented as mean ± standard error of the mean (SEM). Differences were considered significant when p-value is less than 0.05 (* p ≤ 0.05, ** p ≤ 0.01, *** p ≤ 0.001, **** p ≤ 0.0001). Plots and statistical analysis were produced using GraphPad Prism 7 software.

## Acknowledgments

This work was supported by a grant of the European Research Council (ERC, grant N. 322749) to GC.

## Contributions

Experiments has been designed, performed and analysed by GG, SV, GM and FR. Sorting strategy has been designed by GG, EG and CF. Analysis of the Tabula Muris Senis dataset has been carried out by AP. CF, AR performed mass cytometry experiments and GG analysed the output. Project has been supervised by CF, CG, MV, LC and GC. GG and GC wrote the manuscript. All authors reviewed the manuscript.

