## Supplementary Material for "SCA-1 micro-heterogeneity in the fate decision of dystrophic fibro/adipogenic progenitors"

**Supplementary Materials**

**Analysis of the antigen expression data from the multiplex flow cytometry assay**

In order to identify FAPs among the mononuclear cell populations of the skeletal muscle, we reduced the dimensionality of our data by applying the t-distributed stochastic neighbour embedding (tSNE) algorithm. Density plots of the resulting viSNE maps highlight a redistribution of the abundance of cell populations in *mdx* mice when compared to wild type (Supplementary Figure 1A), suggesting a modulation of the profile of muscle mononuclear cell populations in the two mouse models. Next, we applied the FlowSOM algorithm to identify 25 metaclusters of cells with distinct expression profiles and we mapped these metaclusters onto the viSNE maps to facilitate result interpretation (Supplementary Figure 1B). Finally, we scrutinized the antigens expressed in the different metaclusters by consulting the viSNE maps (Supplementary Figure 1C) and we collapsed the metaclusters, showing comparable antigen expression patterns, into new larger clusters.

**Supplementary figure legends**

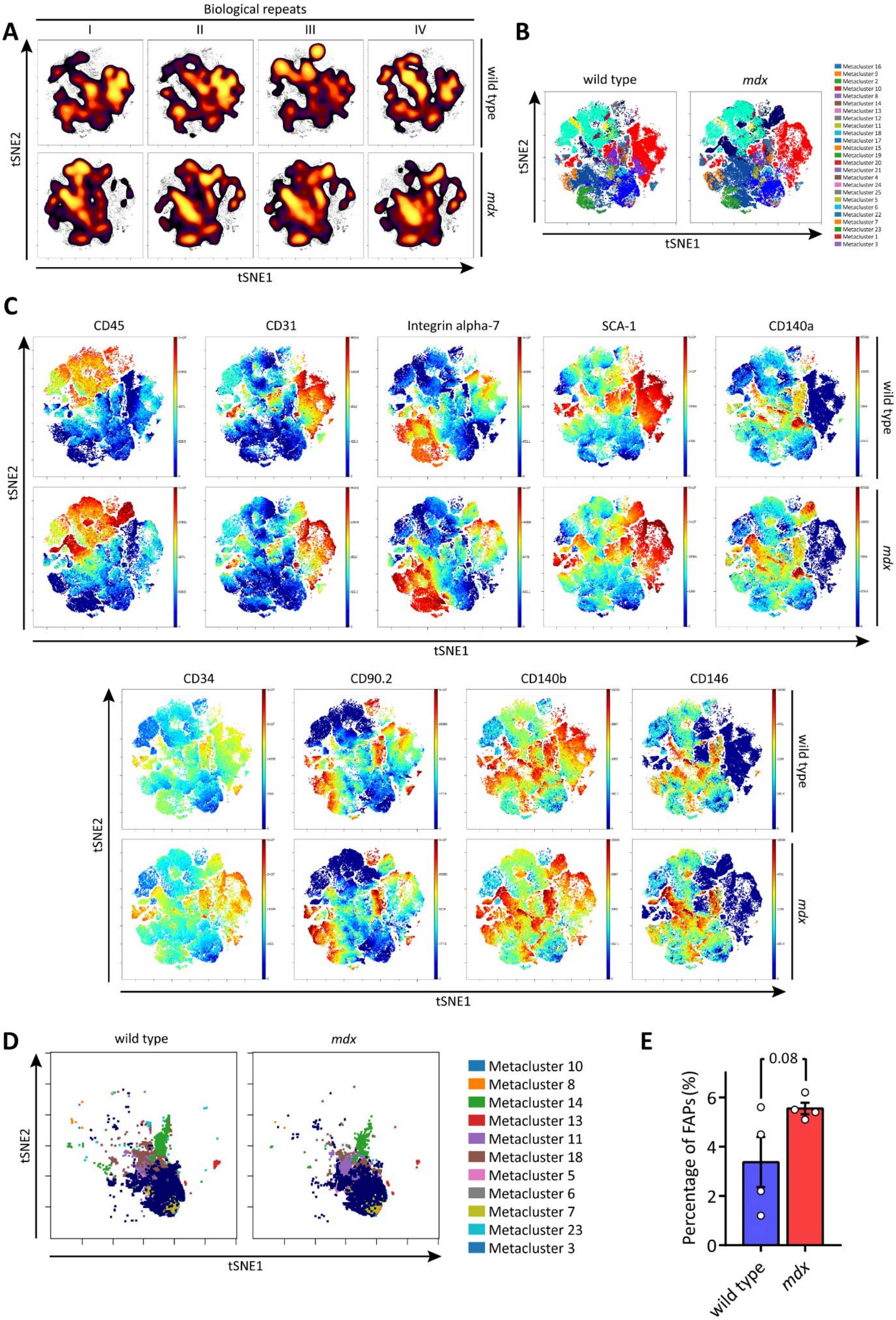

A
Biological repeats
I
II
III
IV
wild type
mdx
tSNE2
tSNE1
B
wild type
mdx
tSNE2
tSNE1
Metacluster 16
Metacluster 9
Metacluster 2
Metacluster 10
Metacluster 8
Metacluster 14
Metacluster 13
Metacluster 12
Metacluster 11
Metacluster 18
Metacluster 17
Metacluster 15
Metacluster 19
Metacluster 20
Metacluster 21
Metacluster 4
Metacluster 24
Metacluster 25
Metacluster 5
Metacluster 6
Metacluster 22
Metacluster 7
Metacluster 23
Metacluster 1
Metacluster 3
C
CD45
CD31
Integrin alpha-7
SCA-1
CD140a
wild type
mdx
tSNE2
tSNE1
CD34
CD90.2
CD140b
CD146
wild type
mdx
tSNE2
tSNE1
D
wild type
mdx
tSNE2
tSNE1
Metacluster 10
Metacluster 8
Metacluster 14
Metacluster 13
Metacluster 11
Metacluster 18
Metacluster 5
Metacluster 6
Metacluster 7
Metacluster 23
Metacluster 3
E
0.08
Percentage of FAPs (%)
0
2
4
6
wild type
mdx

**Supplementary Figure 1. Identification of fibro/adipogenic progenitors among the total mononuclear cells.**

(A) Density representation of viSNE maps showing a redistribution of density patterns between wild type and *mdx* mice. Colour scale ranges from black (low density) to yellow (high density). Each plot represents a biological replicate. (B) Representative viSNE maps illustrating the 25 metaclusters created by the FlowSOM algorithm in wild type and *mdx* mice. These metaclusters were mapped onto the viSNE maps to facilitate the interpretation of the results. Each metacluster is indicated by a different colour. (C) viSNE maps showing the expressing of antigens used in the clustering. These viSNE maps were used to identify metaclusters having similar expression profiles (this is the expert manual refinement phase of our method). The result of such approach are the four clusters in Figure 1B (the four main muscle populations) and 11 unassigned clusters in Supplementary Figure 1D. (D) Representative viSNE maps showing the 11 unassigned clusters. (E) Bar plot indicating the percentage of fibro/adipogenic progenitors (FAPs) in wild type and *mdx* mice. Data are presented as mean ± SEM. Statistical significance was estimated by a Student t-test, * $p \leq 0.05$. ($n = 4$)

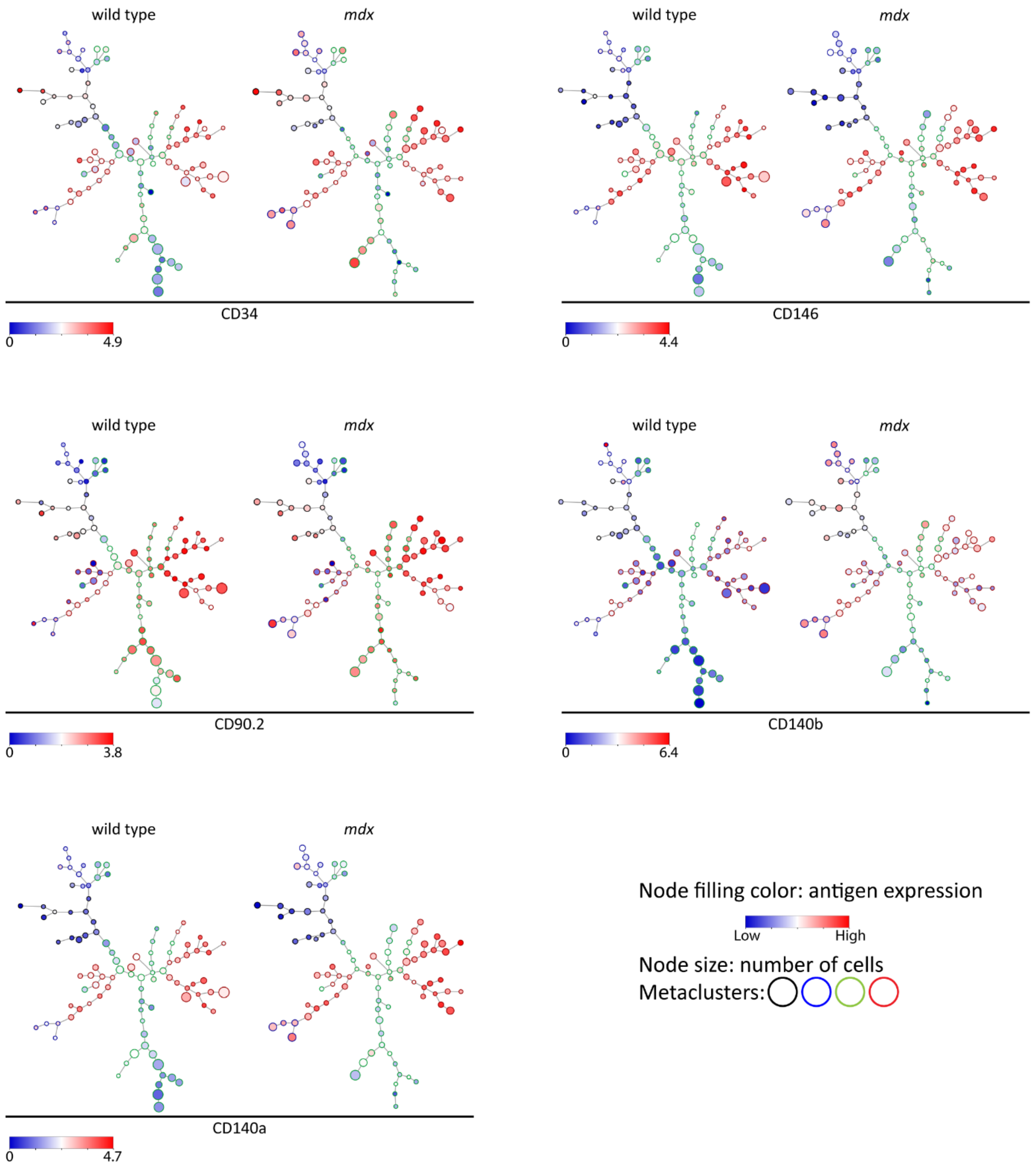


**Supplementary figure 2. Self-organising maps (SOMs) representing expression variability of surface antigens in fibro/adipogenic progenitors.**

The expression of CD34, CD146, CD90.2, CD140b, CD140a and SCA-1 in FAPs (identified through the approach described in Supplementary Figure 1) were analysed by the FlowSOM algorithm. FlowSOM output are the self organising maps (SOMs) showed in this Supplementary Figure. Each node represents a group of cells (clusters). Nodes with similar expression patterns were grouped into 4 metaclusters indicated by their outline colour: black, blue, green and red. Node size is proportional to the number of cells in each cluster. Node filling colour represents the expression level of the antigen ranging from low level (blue) to high level (red). SOM showing SCA-1 expression is reported in Figure 1C.

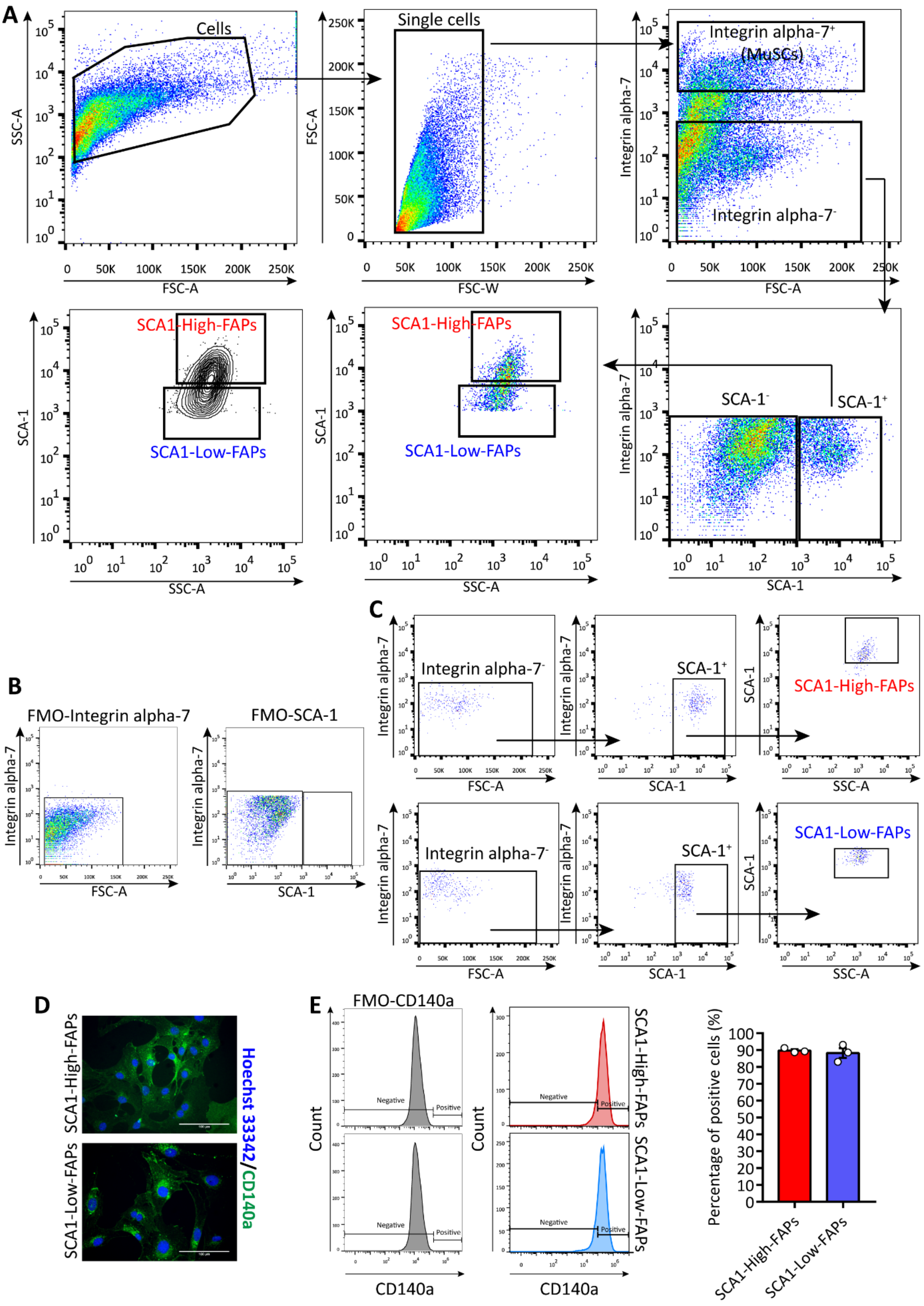


**Supplementary Figure 3. Sorting strategy to isolate fibro/adipogenic progenitors from *mdx* mice.**

(A) Representative dot plots showing the sorting strategy to isolate FAP cell states from hind limb muscles of *mdx* mice. $CD45^-$ $CD31^-$ cells were isolated by magnetic activated cell sorting (MACS). Next, FAP cell states were isolated by fluorescent activated cell sorting (FACS) as Integrin alpha-$7^-$ SCA-$1^+$ cells. MuSCs were isolated as Integrin alpha-$7^+$ cells. (B) Fluorescence minus one (FMO) controls used to place gates of the sorting strategy. (C) Purity of FAP cell states isolated from *mdx* mice. (D) Representative images of FAP cell states from *mdx* mice immunolabelled for CD140a (green). Nuclei were counterstained with Hoechst 33342 (blue). Scale bar 100 μm. (E) Flow cytometry analysis of CD140a expression in FAP cell states expanded in Cytogrow. From left to right: histograms of FMO controls; histograms of stained cells; bar plot illustrating the percentage of FAP cell states positive for CD140a. Data were presented as mean ± SEM. (n = 3)

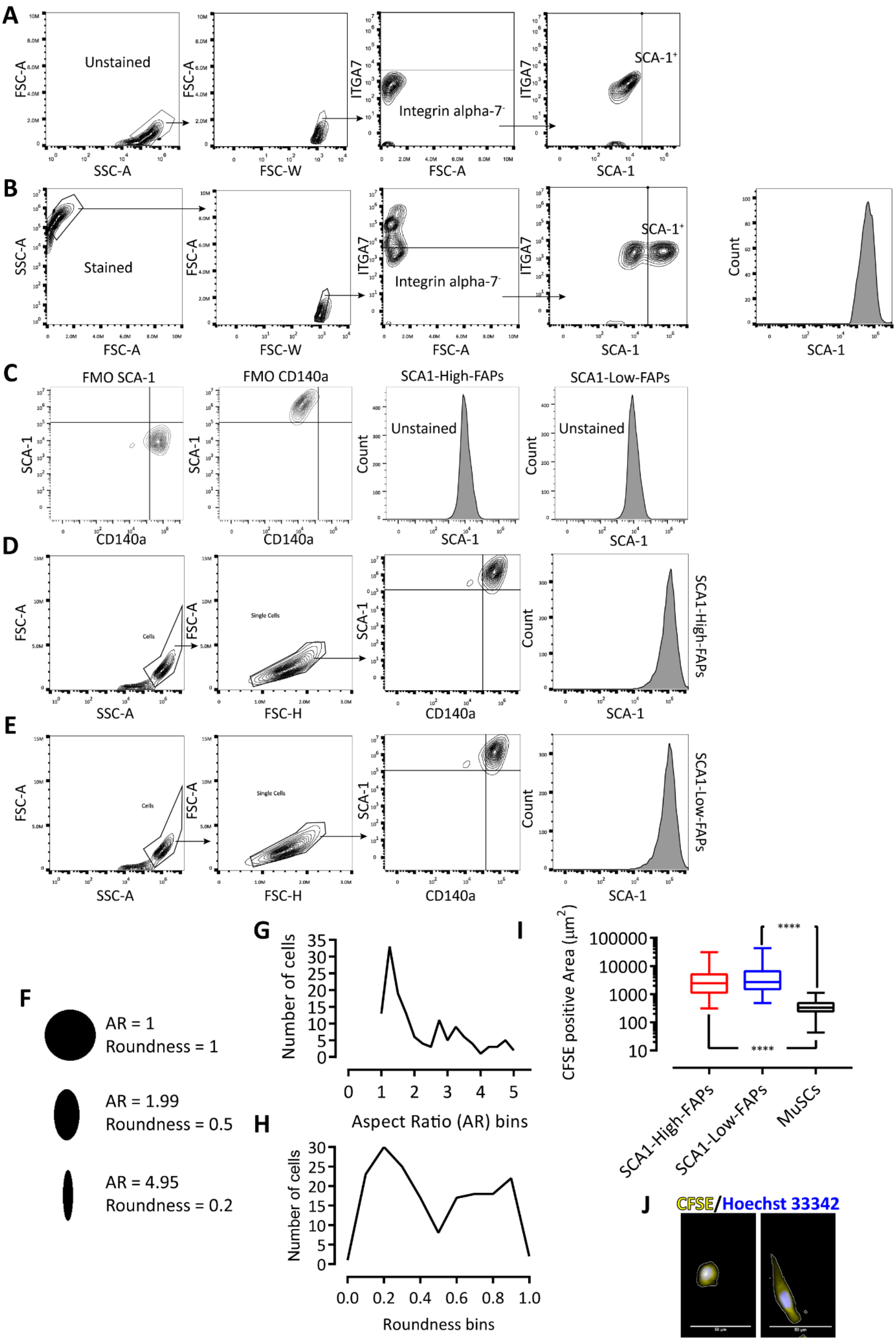
A
Unstained
FSC-A
SSC-A
FSC-W
ITGA7
Integrin alpha-7⁻
FSC-A
SCA-1⁺
SCA-1
B
Stained
SSC-A
FSC-A
FSC-W
ITGA7
Integrin alpha-7⁻
SCA-1⁺
SCA-1
Count
C
FMO SCA-1
FMO CD140a
SCA1-High-FAPs
SCA1-Low-FAPs
Unstained
SCA-1
CD140a
Count
D
FSC-A
SSC-A
Cells
Single Cells
FSC-H
SCA-1
CD140a
Count
SCA1-High-FAPs
E
FSC-A
SSC-A
Cells
Single Cells
FSC-H
SCA-1
CD140a
Count
SCA1-Low-FAPs
F
AR = 1
Roundness = 1
AR = 1.99
Roundness = 0.5
AR = 4.95
Roundness = 0.2
G
Number of cells
Aspect Ratio (AR) bins
H
Number of cells
Roundness bins
I
CFSE positive Area (μm²)
****
SCA1-High-FAPs
SCA1-Low-FAPs
MuSCs
J
CFSE/Hoechst 33342

**Supplementary figure 4. SCA1-High FAPs and SCA1-Low-FAPs are two dynamic cell states and share the same morphology.**

(A) and (B) report representative gating strategy applied to obtain SCA-1 histogram of FAPs in Figure 2A. In (A) are reported contour plots of unstained $CD45^-$ $CD31^-$ cells. In (B) are reported contour plots of $CD45^-$ $CD31^-$ cells stained with antibodies against Integrin alpha-7 and SCA-1. $CD45^-$ $CD31^-$ cells were isolated by MACS from hind limb muscles of *mdx* mice. (C) FMO controls and unstained cells of samples in (D), (E) and Figure 2A. (D) and (E) show gating strategies to study SCA-1 expression in SCA1-High-FAPs and SCA1-Low-FAPs, respectively. Moreover, here we demonstrate that these cells express also CD140a. (F) Aspect ratio (AR) and roundness values changing from a round shape to an elongated shape. (G) and (H) are the distributions of AR and roundness values in *mdx* MuSCs (n = 181) cultured *ex vivo* and stained using CFSE. These values derived from three different biological replicates. Distributions highlight different peaks corresponding to round myoblasts (high roundness/ low AR) and elongated myocytes (low roundness/ high AR). (I) Box plots reporting the CFSE positive area expressed in $\mu m^2$ of SCA1-High-FAPs (n = 96), SCA1-Low-FAPs (n = 94) and MuSCs (n = 181). Whiskers are minimum and maximum values of the distributions. (J) Crop of representative images of round (left) and elongated (right) *mdx* MuSCs stained with CFSE (yellow) and counterstained with Hoechst 33342 (blue). Scale bar 50 $\mu$m. Statistical significance was estimated by a two tailed Mann-Whitney test after a normality test. **** $p < 0.0001$

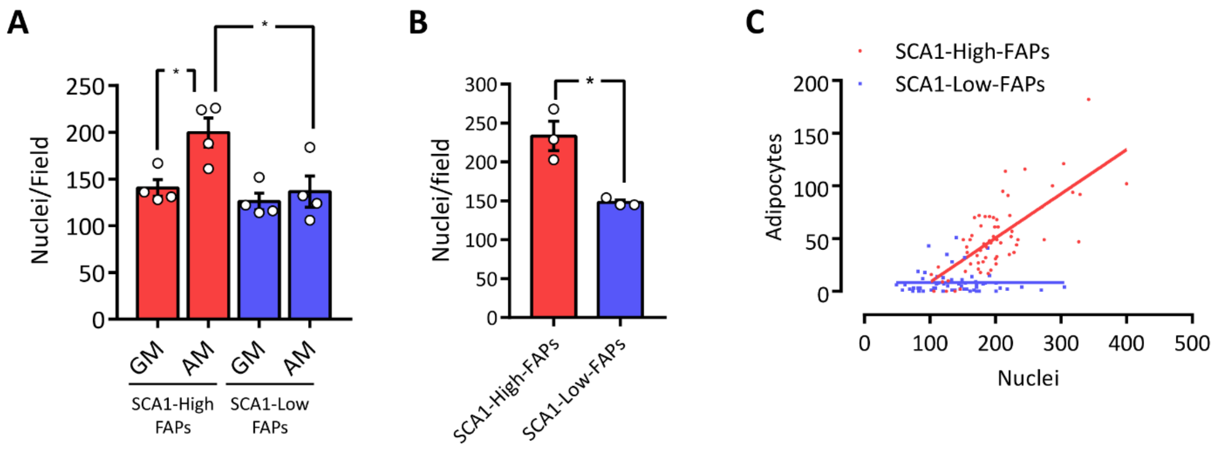


**Supplementary figure 5. Nuclei quantification of FAP cell states induced to differentiate into adipocytes.**

(A) Bar plot reporting nuclei quantification per filed of the experiment in Figure 3B-C (n = 4). (B) Bar plot representing the number of nuclei per filed of the experiment in Figure 3F-G (n = 3). Statistical analysis was performed using Two-way ANOVA in (A) and two tailed Student t-test in (B). (C) Dot plot representing the number of nuclei over the number of adipocytes in each field for SCA1-High-FAPs (in red) and SCA1-Low-FAPs (in blue) from the experiment in Figure 1A-D. Trend lines were calculated using GraphPad Prism 7 software. Data are presented as mean ± SEM. * p< 0.05.

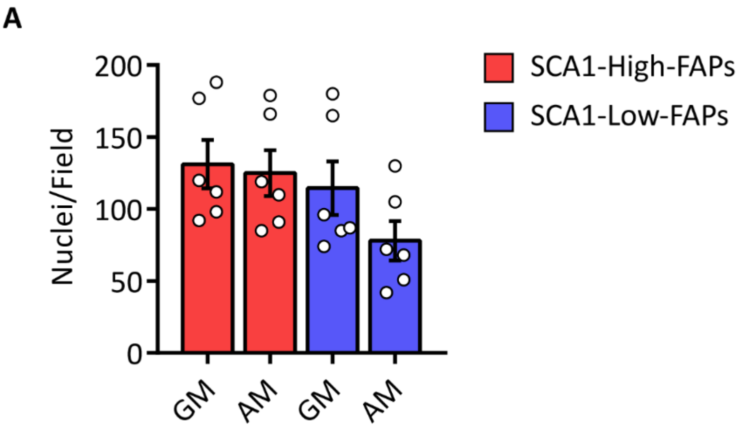


**Supplementary figure 6. Nuclei quantification of FAP cell states induced to differentiate into myofibroblasts.**

(A) Bar plot of nuclei per field of the experiment in Figure 4B (n = 6). Statistical analysis was evaluated using Two-way ANOVA. Data are presented as mean ± SEM

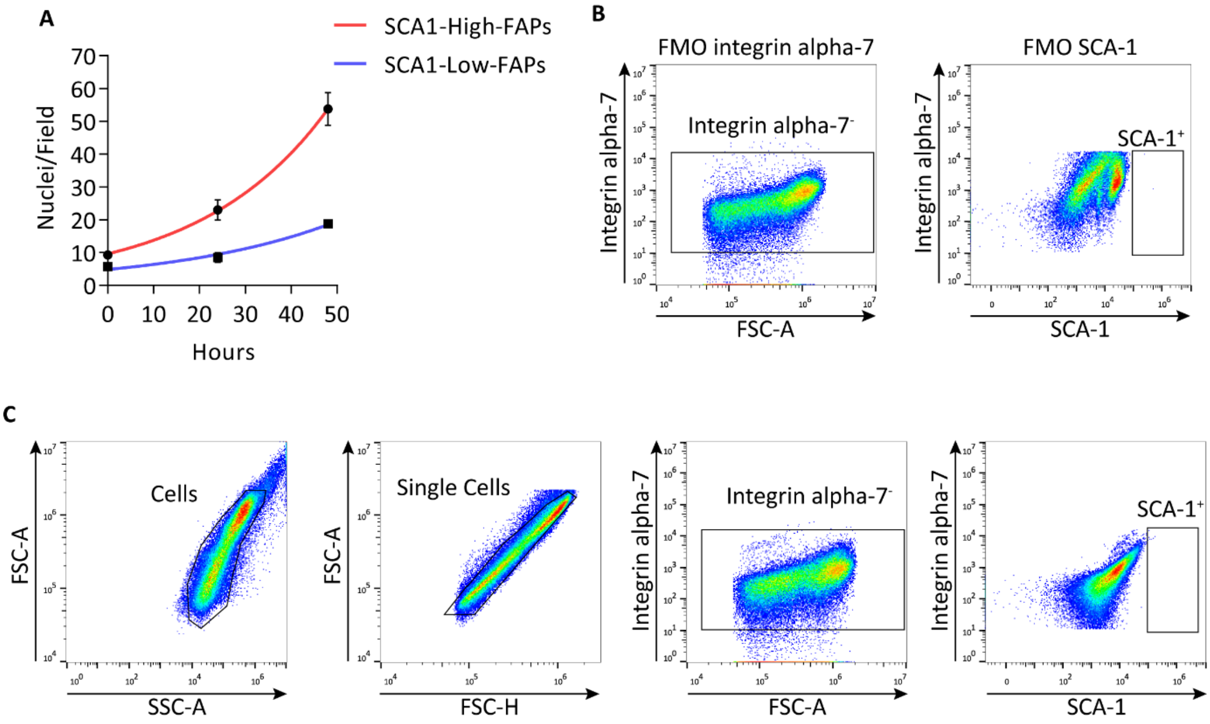


**Supplementary figure 7.** ***In vitro*** **and** ***in vivo*** **proliferation of FAP cell states.**

(A) Nonlinear regression curves of T0, 24 h and 48 h time points of the growth curves in Figure 5B. This curves and doubling times were calculated using the Exponential growth equation tool of Graph Pad Prism. Data are presented as mean ± SEM. (n = 4). (B) and (C) are respectively FMO controls and unstained cells of samples in Figure 5G.

A

| Cell types | Markers |
|---|---|
| Endothelial cells | Gplhbp1, Cdh5, Esam |
| Endothelial cells | Ly6a,Cdh5,Kdr, Pecam1 |
| FAPs | Pdfgra, Ly6a, Vlm, Cd34 |
| Macrophages | Cd68, Ptprc, Itgam, Cd14 |
| Myocytes | Atp2a1, Tcap, Myoz1, Actc1 |
| Neurons | Sfrp5, Kcna1, Matn2 |
| MuSCs | Itga7, Myod1, Vcam1, Sdc4 |
| Smooth muscle cells | Rgs5, Myl9, Kcnj8, Vtn |
| T-cells | Cd3d, Cd3e, Cd3g, Cd4 |
| Tenocytes | Scx, Tnmd, Col1a1, Tnc |
| B-cells | Cd19, Cd79a, Cd22, Ms4a1 |

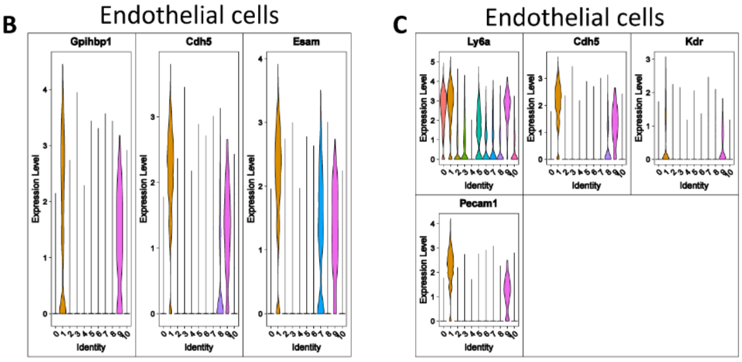


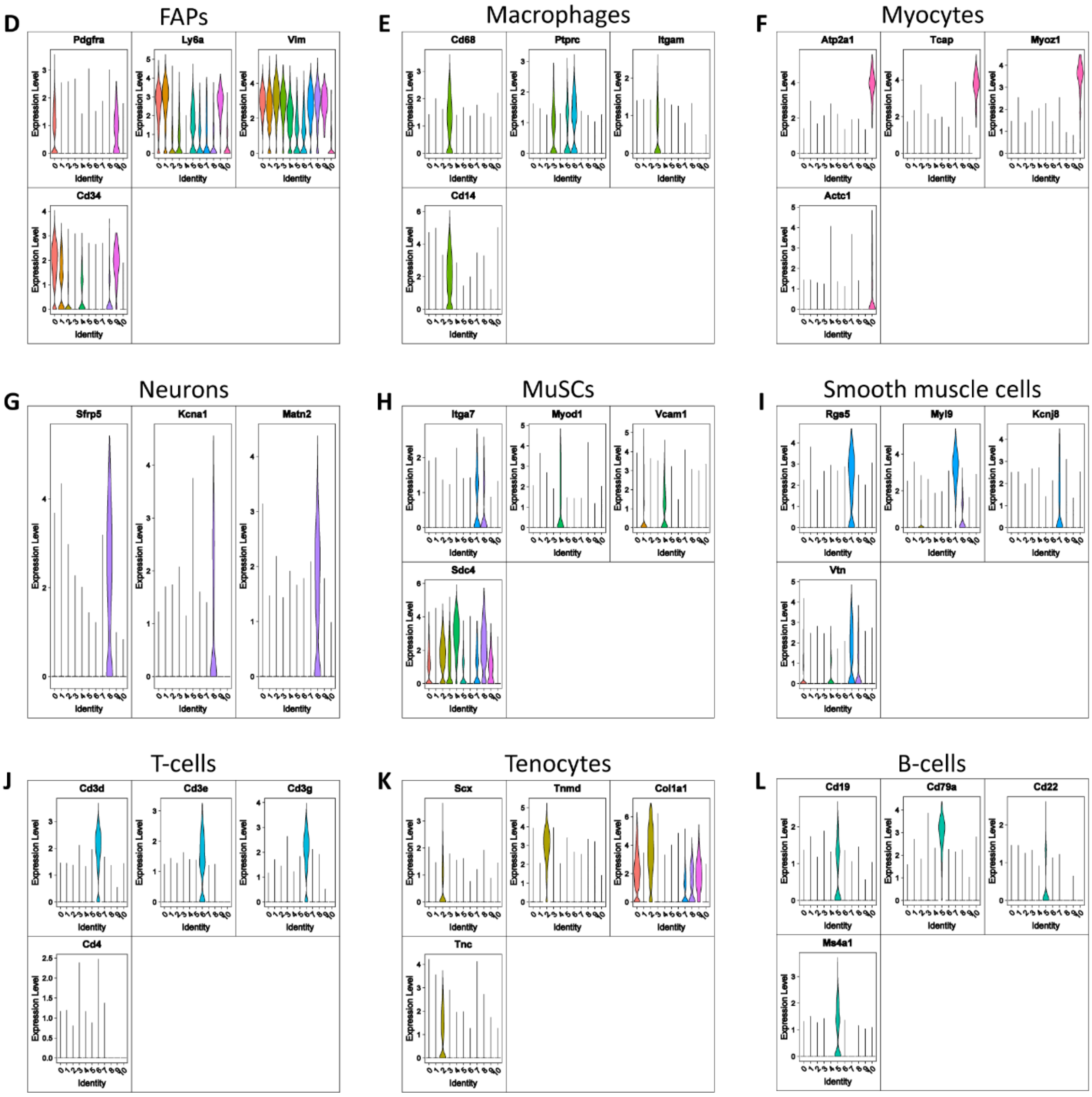

**Supplementary figure 8. Analysis at single-cell level of FAP cell states in wild type mice taking advantage of the Tabula Muris Senis dataset.**

(A) Table reporting the biomarkers used to assign a cell line ontology ID to each cluster. (B-L) Violin plots displaying the distribution of each biomarker across the different clusters for all the identified cell populations.
